# Single-cell lineage tracing identifies hemogenic endothelial cells in the adult mouse bone marrow

**DOI:** 10.1101/2025.10.09.681472

**Authors:** Jing-Xin Feng, Mei-Ting Yang, Lili Li, Caiyi C. Li, Ferenc Livák, Jack Chen, Yongmei Zhao, Dunrui Wang, Avinash Bhandoola, Naomi Taylor, Giovanna Tosato

## Abstract

During mouse development, hematopoietic stem and progenitor cells (HSPC) originate from hemogenic endothelial cells (ECs) through a process of endothelial-to-hematopoietic transition. These HSPC are thought to fully sustain adult hematopoiesis. However, it remains unknown whether adult ECs retain hemogenic potential. Here we used in vivo genetic lineage tracking at population and single-cell (sc) levels, scRNA sequencing, and bone marrow transplantation to detect hemogenic ECs in adult mice. We identify and characterize bone marrow-resident, adult *Cdh5*/VE-Cadherin^+^ ECs that produce hematopoietic cell-progeny in vitro and in mice. These adult hemogenic ECs and their hematopoietic cell progeny give rise to hematopoietic cells following adoptive transfer into adult mice. Furthermore, blood cells generated from adult and developmental ECs comparably home to peripheral tissues, where they similarly contribute to inflammatory responses. Thus, our results identify previously unrecognized bone marrow-derived adult hemogenic ECs that generate HSPC and functional mature blood cells.

## INTRODUCTION

During development, hemogenic ECs generate hematopoietic cells through a process of endothelial-to-hematopoietic transition (EHT) at geographically defined anatomical sites^1–9^. At these locations, the hemogenic ECs represent a small fraction of all ECs^10–12^, and their competency to produce hematopoietic stem and progenitor cells (HSPC) is temporally restricted to short developmental windows, and the hemogenic potential differs^13–18^. In the dorsal aorta, the hemogenic endothelium produces hematopoietic stem cells (HSC) and other multi-potent progenitors between E10.5 and E11.5^17–19^ that persist in the adult and are generally believed to sustain adult hematopoiesis throughout life.

Efforts to reliably generate *ex vivo* HSC from Cdh5-expressing ECs identified difficulties in reproducing the microenvironmental clues coming from the inductive niche cells^20–22^ and have generally relied on transcription factor-induced reprogramming of the ECs to drive hematopoietic specification^23,24^. The transcription factor RUNX1, which marks the hemogenic endothelium, can confer a hemogenic potential to embryonic ECs lacking such potential^25–28^. When transcription factors RUNX1, FOSB, GFI1, and SPI1 were co-expressed in adult murine ECs co-cultured with “vascular niche ECs”, EHT was induced in vitro producing HSC with long-term self-renewal capacity^29^. A similar approach was used with human ECs enabling hematopoietic specification^30^.

An unresolved question is whether hemogenic ECs, thought to be mostly restricted to the early stages of mouse development^12,23^, may persist in the adult mouse^2,12,31,32^. Exploiting advances in cell lineage tracking and single cell analyses^33,34^, we report the identification of hemogenic ECs in the adult mouse bone marrow (BM) that produce functional hematopoietic progenitors and mature blood cells.

## RESULTS

### Evidence that adult bone marrow ECs generate hematopoietic cells

We assessed the hemogenic potential of ECs in adult mice using Cre-reporter-based lineage tracing. Since *Cdh5*, encoding vascular endothelial cadherin (VE-Cadherin), is selectively expressed by ECs, Cdh5-Cre^ERT2^ recombinase activity can allow tracking the hematopoietic cell output from adult hemogenic ECs^35,36^. Therefore, we generated three Cdh5-based lineage-tracing models using inducible Cdh5-Cre^ERT2^(PAC)^37,38^ and Cdh5-Cre^ERT2^(BAC)^39^ mouse lines, in combination with the Cre-reporter lines ZsGreen and mTmG (Figure S1A). We then treated eight- to twelve-week-old mice with three doses of tamoxifen (10 mg·kg^-1^, gavage) on consecutive days, and four weeks later we evaluated peripheral blood and BM (Figure 1A).

**Figure 1.**
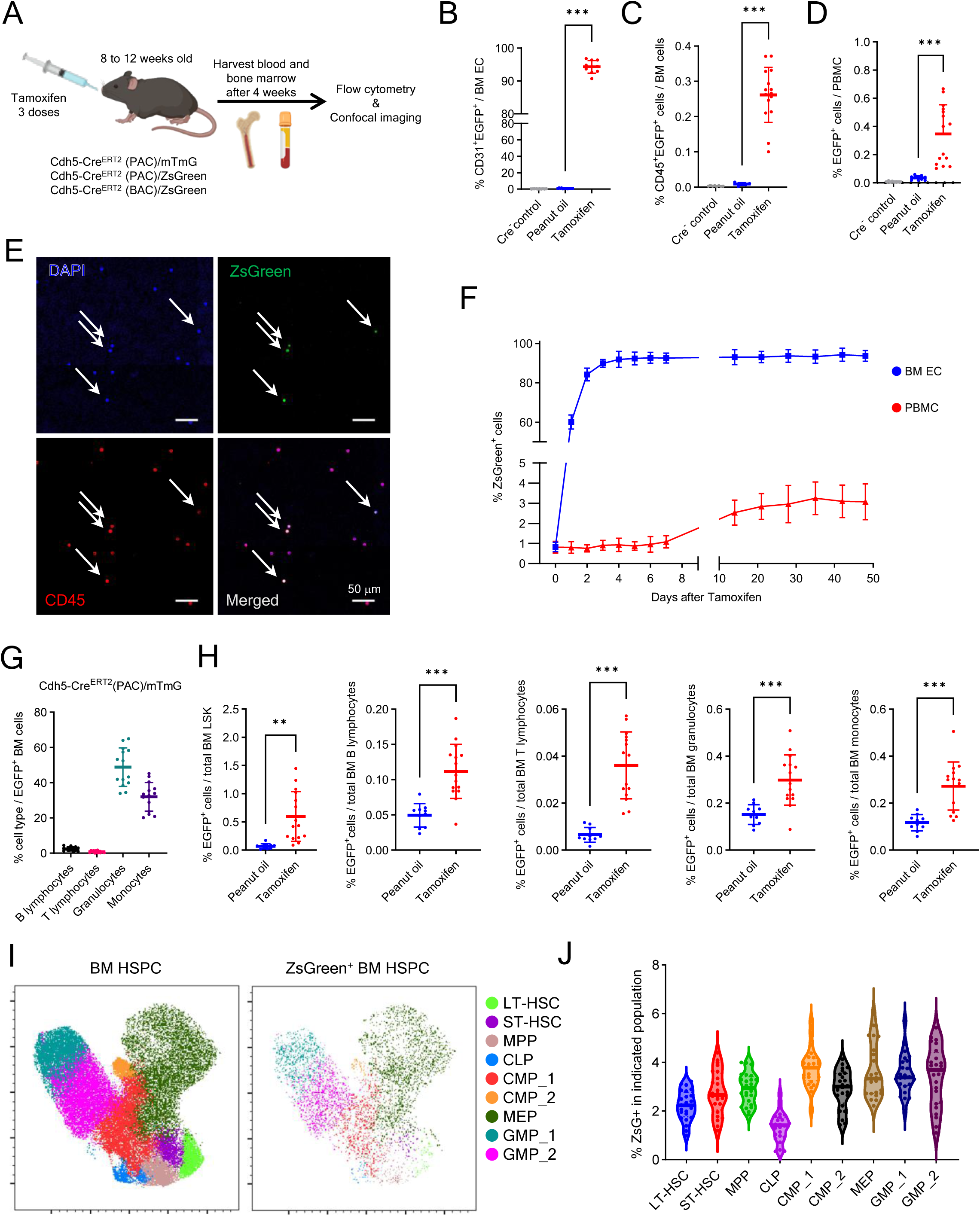
Lineage tracking discloses a contribution of endothelial cells to hematopoiesis in adult BM (A) Experimental design: tamoxifen was administered to 8- to12-week-old Cdh5-Cre mice to induce fluorescent labeling of VE-Cadherin⁺ cells and their cell progeny. Four weeks later, BM and blood were analyzed. (B) CD31⁺EGFP^+^ BM ECs in Cre⁻ mice (n=10) and Cre⁺ mice treated with oil (n=13) or tamoxifen (n=10); flow cytometry results. (C and D) CD45^+^EGFP^+^ cells in BM and blood from Cre⁻ mice (n=8) and Cre⁺ mice treated with oil (n=6-10) or tamoxifen (n=15-18). Representative flow cytometry gating in Figure S1G. (E) Representative blood smear from a tamoxifen-treated Cdh5-Cre^ERT2^(PAC)/ZsGreen mouse showing ZsGreen⁺CD45⁺DAPI⁺ cells (arrows). (F) Kinetics of ZsGreen⁺ cell detection in BM ECs (CD45⁻VE-Cadherin⁺) and blood white blood cells (WBC) post-tamoxifen; mouse n=8-10/group). (G) EGFP^+^ B and T-lymphocytes, granulocytes, and monocytes in BM of tamoxifen-treated mice (n=14) as percent of total EGFP^+^ cells; 3 experiments. (H) EGFP⁺ BM LSK, lymphocytes, granulocytes, and monocytes as percent of total EGFP^⁺/-^ cell type; Cdh5-Cre^ERT2^(PAC)/mTmG mice (oil n=10; tamoxifen n=15), 3 experiments. (I) UMAP plots of Lin⁻ BM HSPC from tamoxifen-treated Cdh5-Cre^ERT2^(PAC)/ZsGreen mice (n=26; 1 femur/mouse) showing FlowSOM clustering of all (ZsGreen⁺/-) and ZsGreen⁺ populations. (J) Violin plots showing ZsGreen⁺ cell distribution across HSPC subsets from (I). Dots represent individual mice; data shown as mean±SD except shown as median in (G). *p < 0.05, **p < 0.01, ***p < 0.001 by Student’s t test.

Expectedly^36^, most BM Endomucin^+^ ECs were ZsGreen^+^ in tamoxifen-treated Cdh5-Cre^ERT2^(PAC)/ZsGreen and Cdh5-Cre^ERT2^(BAC)/ZsGreen mice, and virtually no ZsGreen^+^ ECs were present in Cre negative or peanut oil-treated controls (Figures S1B and S1C). By flow cytometry, >90% CD31^+^VE-Cadherin^+^ BM ECs were tracked by tamoxifen-induced fluorescence in the three mouse lines (Figure 1B; Figure S1D). A low-level tamoxifen-independent reporter fluorescence was also detected in BM ECs, which was low in the mTmG reporter line (Figure 1B), and higher in the ZsGreen reporter lines, previously attributed to “basal” Cre^ERT2^ activity^40^ (Figure S1D).

To evaluate the hemogenic potential of adult BM ECs, we analyzed the expression of the hematopoietic marker CD45 in Cdh5-tracked cells. Notably, tracked CD45^+^ hematopoietic cells, presumed progeny of VE-Cadherin^+^ ECs, were detected by flow cytometry in BM and blood of mice from all three Cdh5-Cre^ERT2^ mouse lines (Figure 1C and D; Figures S1E and F; representative flow cytometry gating in Figure S1G and H). Moreover, confocal microscopy identified isolated ZsGreen^+^CD45^+^ cells in BM (Figure S1I) and blood of tamoxifen-induced mice (Figure 1E). Importantly, while a small fraction of fluorescent CD45^+^ cells were detected in EGFP and ZsGreen (Figures 1C and D; and Figure S1D-F) reporter mice without tamoxifen, as reported^40–43^, the significant increase of tamoxifen-induced fluorescent CD45^+^ cells (Figure 1C and 1D, Figure S1D-F) indicates the occurrence of Cdh5-Cre recombination in the adult mouse, presumably tracking EHT. This tamoxifen-induced increase in CD45^+^ZsGreen^+^ peripheral blood mononuclear cells (PBMC) was also observed in individual mice (Figure S1J).

We tested the kinetics of tamoxifen-induced ZsGreen expression in BM ECs and PBMC of Cdh5-Cre^ERT2^(PAC)/ZsGreen mice (Figure 1F). By flow cytometry, virtually all BM ECs were ZsGreen^+^ by day 4 after tamoxifen administration and this level persisted over 50 days.

Instead, the percentage of ZsGreen⁺ PBMCs increased more gradually, plateauing around day 30 and this level persisted over ∼50 days of observation. This gradual increase is likely attributable to tamoxifen-induced ZsGreen expression in hemogenic ECs.

Further analysis showed that most BM EGFP^+^CD45^+^ cells in Cdh5-Cre^ERT2^(PAC)/mTmG mice were CD11b^+^Ly6G^+^ granulocytes (48.85%) and CD11b^+^Ly6G^-^ monocytes (32.06%), while CD19^+^ B (2.46%) and CD3^+^ T (0.84%) lymphocytes were detected at lower frequencies (Figure 1G). All EGFP^+^ BM cell populations were significantly induced by tamoxifen administration, including rare LSK (Lin^-^Sca1^+^cKit^+^) progenitors (Figure 1H). Also, the peripheral blood of tamoxifen-induced Cdh5-Cre^ERT2^(PAC)/mTmG mice contained EGFP^+^ granulocytes and monocytes, and fewer B and T lymphocytes (Figure S1K and L). Similarly, in ZsGreen-reporter mice, tracked LSK progenitors, B lymphocytes, granulocytes, and monocytes were all present in the BM and blood (Figure S2A-D).

We further characterized the tracked BM hematopoietic LSK stem/progenitors by flow cytometry (Figure S2E). The results, displayed by Uniform Manifold Approximation and Projection (UMAP) dimensional reduction, showed that the ZsGreen-tracked progenitor cell population includes phenotypic subsets consistent with hematopoietic stem and progenitor cells (HSPC) (Figure 1I). Each ZsGreen^+^ progenitor population represented a similarly small proportion of the corresponding non-tracked progenitor cell population (Figure 1J). These results support the existence of EHT in the adult mouse contributing to generation of hematopoietic progenitors and mature cells.

### Adult BM ECs cultured ex vivo generate transplantable HSPCs

To investigate whether adult BM ECs can generate hematopoietic cells *ex vivo*, we cultured BM cells isolated from tamoxifen-treated Cdh5-Cre/ZsGreen mice. Initially, BM single-cell (sc) suspensions were cultured under two conditions (Figure 2A): (1) on “Primaria” pro-adhesive flasks, and (2) on OP9 stromal cell^44^ monolayers grown on gelatin-coated conventional tissue culture flasks. To attempt recreating BM niches, fresh wild-type (WT) BM cells were added to the ZsGreen-tracked BM cell cultures twice/week throughout the culture period (see Methods for details). By week 8, BM cells cultured on “Primaria” dishes formed a confluent monolayer of ZsGreen-tracked cells with a typical EC morphology, whereas BM cells cultured on OP9 monolayers exhibited a fibroblast-like morphology with the ZsGreen^+^ cells clustering in foci, without forming a monolayer (Figure 2A). We visualized some round ZsGreen-tracked (ZsGreen^+^) cells in cultures of BM cells grown onto OP9 monolayers supplemented with fresh BM cells, but not in cultures of BM cells grown onto “Primaria” surfaces supplemented with fresh BM cells (Figure S2F). This hinted that the OP9 and BM cell culture system may allow hematopoietic cells emergence from Cdh5-Cre/ZsGreen-tracked EC ex vivo.

**Figure 2.**
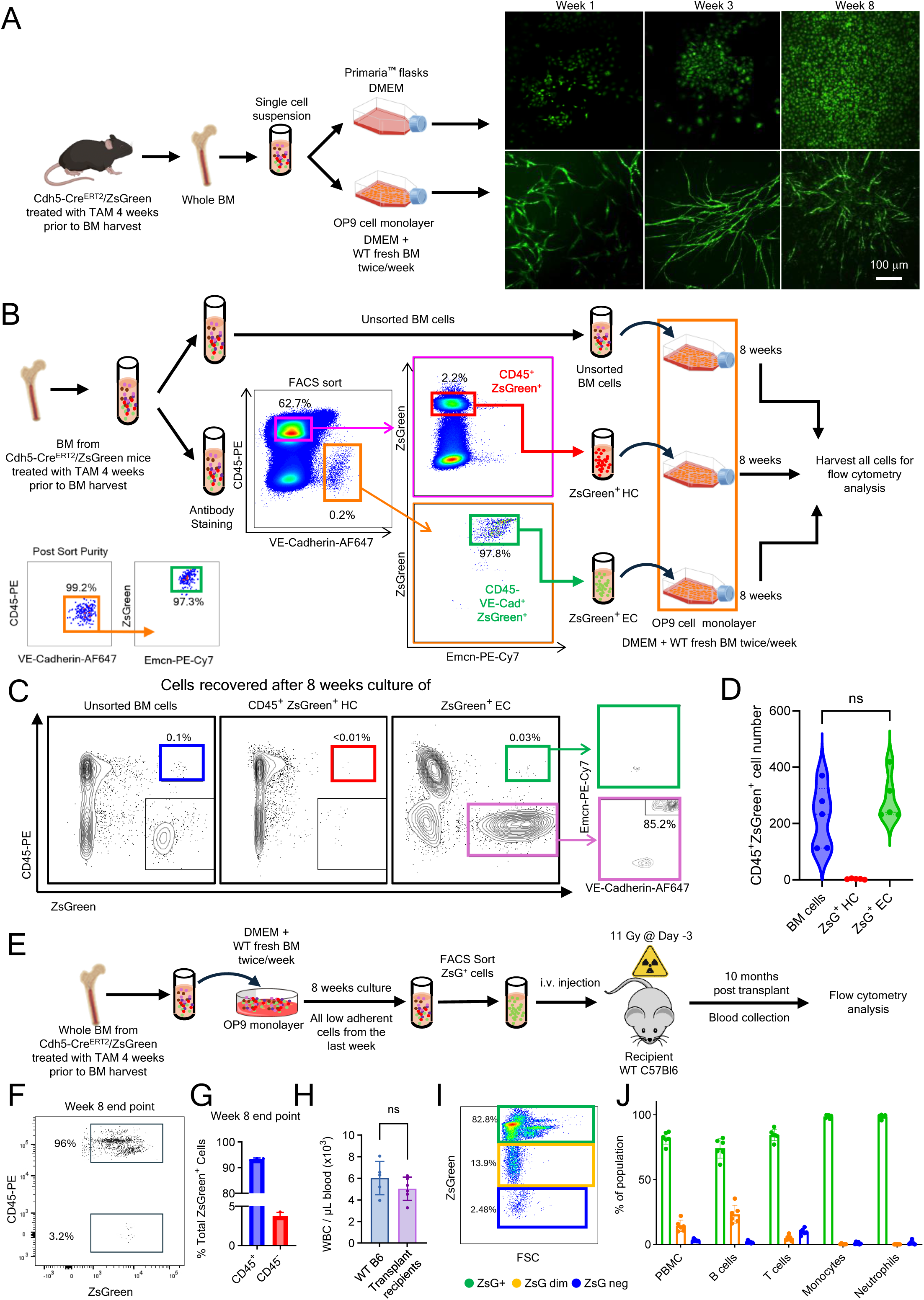
BM ECs generate engraftable hematopoietic cells ex vivo (A) BM cells from tamoxifen-treated mice were cultured on high-attachment Primaria flasks or OP9 cell monolayers. Representative images show ZsGreen⁺ cells at weeks 1, 3, and 8. (B) Workflow for culturing unsorted and sorted BM cell populations. Post-sort purity of ZsGreen^+^ ECs is shown in the bottom left panel. All cells were cultured (8 weeks) on OP9 cell monolayers supplemented with WT BM cells. Culture medium and floating cells were removed twice/week for 7 weeks. At the start of week 8, one final WT BM and medium supplementation was implemented prior to harvest at the end of week 8. (C and D) Representative flow cytometry plots (C) and quantification (D) of CD45⁺ZsGreen⁺ cells from each of the 8-week cultures (n=5). (E) Floating/loosely adherent ZsGreen⁺ cells from unsorted BM 8-week cell cultures were sorted and transplanted (5×10^4^, 2.5×10^4^,1.25×10^4^ or 6.25×10^3^ cells) into lethally irradiated (11 Gy) WT mice (n=2/group). (F and G) Representative flow cytometry image (F) and quantification (G) of low-adherent cells harvested after 8 weeks of culture, showing that >95% (group average) of ZsGreen⁺ low-adherent cells are CD45⁺. These ZsGreen^+^CD45^+^ cells were sorted for transplantation. (H - J) WBC counts from 5 control mice (no irradiation or transplant) (H) and percent ZsGreen⁺, ZsGreen dim and ZsGreen⁻ cells (I-J) in blood of transplant recipients 10 months post-transplant (n=6). Dots represent individual mice. Data are shown as mean ± SD. ns, not significant by Student’s *t* test.

To test this possibility, we generated a pool of BM cells (from 20 adult mice; age 10-18 weeks) and used the pool to derive 3 populations: sorted BM CD45^−^VE-Cadherin^+^ZsGreen^+^ ECs; sorted CD45^+^ZsGreen^+^ hematopoietic cells; and unsorted BM cell populations (Figure 2B). We then cultured these 3 cell populations under the same conditions (OP9 monolater and supplementation with fresh BM cells twice/week). After 8-week culture, flow cytometry analysis detected CD45^+^ZsGreen^+^ cells from the unsorted BM and the sorted CD45^−^VE-Cadherin^+^ZsGreen^+^ cell cultures. However, CD45^+^ZsGreen^+^ were virtually absent from the CD45^+^ZsGreen^+^ cell cultures (Figure 2C and 2D), likely attributable to the culture conditions not designed to support growth, differentiation or survival of hematopoietic cells. These results show that CD45^−^VE-Cadherin^+^ZsGreen^+^ ECs can generate CD45^+^ZsGreen^+^ *ex vivo*. Notably, most (>80%) CD45⁻ZsGreen⁺ cells retained expression of VE-Cadherin and Endomucin, thereby confirming their endothelial identity (Figure 2C, most right panel). Importantly, the virtual absence of CD45^+^ZsGreen^+^ cells in 8-week cultures of sorted CD45^+^ZsGreen^+^ cells shows that pre-existing CD45^+^ZsGreen^+^ hematopoietic cells, derived from tamoxifen-dependent and independent processes, are effectively removed during the extended culture. This further suggests that CD45^+^ZsGreen^+^ hematopoietic cells, potentially contaminating the sorted ECs, are unlikely contributors to EC-derived hematopoiesis. Rather, these results show that the ECs are the likely cell source of hematopoietic cells in this *ex vivo* co-culture model.

Next, we tested whether the *ex vivo*-derived CD45^+^ZsGreen^+^ cells are functional in lethally irradiated recipients (Figure 2E). To this end, we obtained unfractionated BM cells from 50 tamoxifen-treated Cdh5-Cre/ZsGreen mice, cultured the cells onto OP9 monolayers with fresh BM supplementation, sorted the ZsGreen^+^ cells (>95% of which were CD45^+^ by flow cytometry) at the end of 8-week culture, and transplanted these cells into 8 lethally irradiated WT recipients; 5×10^3^ cells; 2.5×10^3^ cells; 1.25×10^3^ cells; 6.25×10^2^ cells, n=2/group (Figure 2E-G). Two mice (recipients of 6.25×10^2^ and 1.25×10^3^ cells) died on days 4 and 6, but the remaining six mice survived and remain well at the time of manuscript revision (10 months post-transplant).

Ten months post-transplant, peripheral blood WBC counts were within normal range in all surviving transplant recipients, indicative of hematopoietic reconstitution (Figure 2H). Flow cytometry revealed that 81.9% of PBMC were ZsGreen^+^ (Figure 2I-J). Most neutrophils (98.6%), monocytes (98.5%), B cells (74.1%) and T cells (84.3%) were ZsGreen^+^. These results confirm that BM ECs propagated *ex vivo* onto OP9 monolayers with BM cell supplementation produce hematopoietic cells and show that this output includes HSPC capable of engrafting and generating hematopoietic cell progeny.

### Adult BM ECs can give rise to hematopoietic cells following transfer into conditioned recipients

Next, we examined whether adult BM ECs are hemogenic in transplant recipients. To this end, we FACS-sorted CD45^-^VE-Cadherin^+^ZsGreen^+^ ECs from the BM of tamoxifen-pretreated (4 weeks prior to BM harvest) Cdh5-Cre^ERT2^(PAC)/ZsGreen mice (Figure 3A; Figure S2G and S2H) and transplanted these cells into adult WT C57Bl/6 unconditioned mice (PBS) or conditioned by fluorouracil (5-FU treated) prior to transplant^45^. The choice of 5-FU conditioning rather than lethal irradiation of adult mice was driven by previous experiments showing the difficulties at reconstituting lethally myelo-ablated adult recipients with hemogenic yolk sac cells, which reconstituted conditioned newborns^46^. Four weeks after transplant, we observed a significant increase in the proportion of ZsGreen^+^ ECs in the BM and ZsGreen^+^CD45^+^hematopoietic cells in the BM and peripheral blood of the transplant recipients conditioned by 5-FU but not controls (PBS) (Figure 3B and 3C; Figure S2I). These CD45^+^ZsGreen-tracked cells in BM and blood included granulocytes, monocytes, and lymphocytes (Figure 3D). Thus, BM-derived CD45^-^VE-Cadherin^+^ZsGreen^+^ cells transferred into 5-FU-conditioned recipients gave rise to detectable CD45^+^ZsGreen^+^ hematopoietic cells.

**Figure 3.**
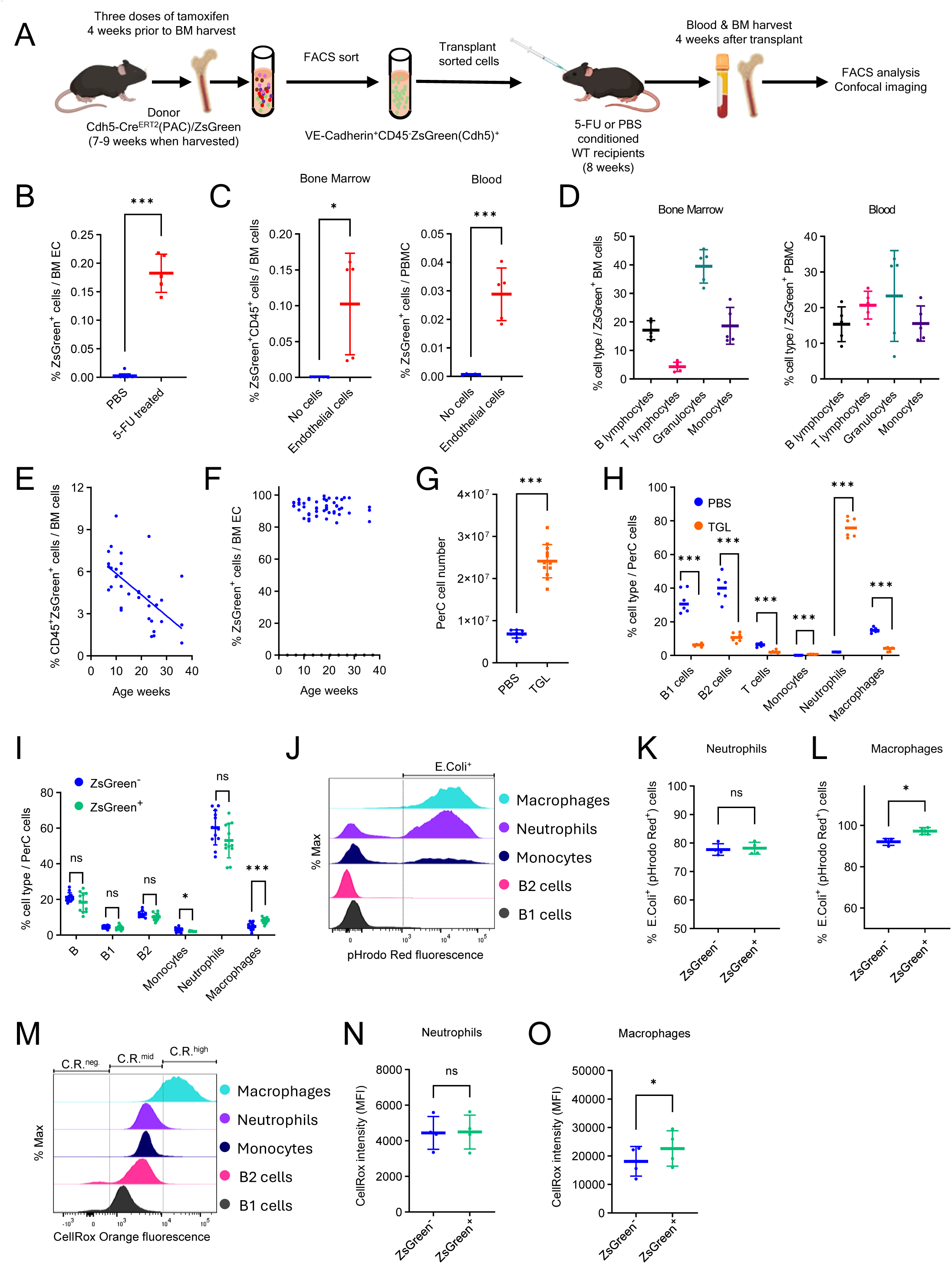
Adult BM endothelial cells give rise to hematopoietic cells following transfer into conditioned recipients (A) Transplant experiment: donor ECs from BM of tamoxifen-treated mice were FACS-sorted and transplanted into WT C57Bl/6 recipients conditioned with 5-FU or PBS. (B) ZsGreen⁺ ECs detected in BM of 5-FU-conditioned (n=5) or PBS-conditioned (n=15) recipients of ECs 4 weeks post-transplant. (C and D) ZsGreen^+^CD45^+^ cells (C) and cell type distribution (D) in the BM and blood of 5-FU-conditioned transplant recipients of BM ECs or no cell controls (n=5/group). (E and F) Age-dependent decline of ZsGreen⁺CD45⁺ cells (E) but not ZsGreen^+^VE-Cadhenin^+^ cells (F) in the BM of Cdh5-Cre^ERT2^(BAC)/ZsGreen mice (n=35) treated with tamoxifen 4-weeks prior to harvest. (G and H) Cell number (G; mouse n=8-12) and cell type distribution (H; mouse n=6) in the peritoneal cavity (PerC) of PBS- or thioglycolate (TGL)-pretreated (4 hours) mice. (I) ZsGreen⁺ and ZsGreen⁻ PerC cell types in TGL-pretreated mice (n=12). (J) Representative histograms depicting pHrodo Red fluorescence detection of E-Coli phagocytosis. (K and L) E. coli⁺ phagocytosis by ZsGreen⁺ and ZsGreen⁻ PerC neutrophils (K) and macrophages (L) in TGL-pretreated mice (n=4). (M) Representative histograms depicting CellRox Orange fluorescence for cell-associated ROS detection. (N and O) CellRox mean fluorescence intensity (MFI) in ZsGreen⁺ and ZsGreen⁻ PerC neutrophils (N) and macrophages (O) in TGL-pretreated mice (n=4). Dots represent individual mice. Data are shown as mean ± SD. *p < 0.05, ***p < 0.001, ns, not significant by Student’s *t* test.

We further examined the effect of mouse age on the endothelial hemogenic potential by treating the mice with tamoxifen between week 6 and 32 of age. Four weeks later (week 10 to 36 of age), we measured the percentage of ZsGreen-tracked CD45^+^ hematopoietic cells in the BM. We observed that tamoxifen inductions beyond week 10 of age resulted in a progressive decrease of the CD45^+^ hematopoietic cell output, and detected an inverse correlation (Pearson’s r −0.63, *P*<0.0001) between age and EC hemogenic potential (Figure 3E). This progressive decline of CD45^+^ cell output was not coupled with a loss of BM EC fluorescence, since virtually all BM ECs were ZsGreen^+^ throughout the duration of the experiment (Figure 3F), consistent with the stability of Cre-mediated labeling of Cdh5-expressing cells ^36^. These observations suggest an age-related loss of hemogenic capability of BM ECs.

Additionally, we examined whether tracked hematopoietic cells from adult BM EC are functional. Since lymphocyte trafficking from the peripheral blood to the peritoneal cavity is critical for their function at this site^47,48^, we first evaluated the spontaneous migration of tracked CD45^+^ hematopoietic cells to the peritoneal cavity. Compared to no-tamoxifen controls, tamoxifen-treated Cdh5-Cre^ERT2^(PAC)/ZsGreen mice displayed a significant increase of CD45^+^ZsGreen^+^ monocytes, macrophages, and B1, B2, and T lymphocytes in the peritoneal cavity (Figure S2J and S2K).

After inducing peritonitis with thioglycolate (TGL, 4 hours) in tamoxifen-treated Cdh5-Cre^ERT2^(PAC)/ZsGreen mice, the overall number of peritoneal leukocytes increased substantially compared to untreated (PBS) controls (Figure 3G), mostly attributable to neutrophils (Figure 3H). In addition, the peritoneal ZsGreen^+^ and ZsGreen^-^ cell populations exhibited a similar cell type distribution in TGL-treated mice (Figure 3I). We further examined *Escherichia coli* (K-12 strain) phagocytosis and reactive oxygen species (ROS) production in peritoneal cell exudates in response to TGL (Figure 3J-3O). Both ZsGreen^+^ and ZsGreen^-^ peritoneal neutrophils and macrophages comparably phagocytosed *Escherichia coli* and generated ROS (Figure 3J, 3K, 3M, and 3N), except that the phagocytic and ROS production of ZsGreen^+^ macrophages was somewhat higher than that of ZsGreen^-^ macrophages (Figure 3L and 3O). Thus, Cdh5-tracked mature neutrophils and macrophages are functional at trafficking and homing, and presumably capable of contributing to the host response to tissue inflammation.

### Adult EHT is independent of preexisting hematopoietic cell progenitors

Faithful Cre-reporter lineage tracing requires that Cre recombinase activity be restricted to the intended cell type^40^. In our system, Cdh5-Cre^ERT2^ is expected to drive recombination specifically in ECs, as *Cdh5* is an established EC-specific marker. However, flow cytometry has occasionally revealed rare VE-Cadherin^+^CD45^+^ in mouse BM, exemplified in Figure S2G, potentially reflecting double-positive cells. To address the possibility that cells co-expressing VE-Cadherin/*Cdh5* and CD45/*Ptprc* exist in the mouse BM, we analyzed publicly available sc-RNAseq data from adult mouse BM^49^. This analysis showed that only a small subset of plasmacytoid dendritic cells (pDCs) co-express VE-Cadherin and CD45, but not HSPC or other mature blood cells (Figure S3A-S3D). pDCs are terminally differentiated cells and unlikely progenitors of tracked CD45^+^ multilineage progeny in our Cdh5-cre mice. Nonetheless, it is plausible that other, currently unidentified, hematopoietic cells may also co-express *Cdh5*/VE-Cadherin and *Ptprc*/CD45 and possess functional Cre^ERT2^ activity, inducing fluorescence in these cells upon tamoxifen administration.

To address these possibilities, we transplanted lethally irradiated (11 Gy) WT C57Bl6 mice (n=6) with ZsGreen⁻Lin⁻Sca1⁺cKit⁺ (LSK) progenitors (>99% purity; Figure S3E-G) from tamoxifen untreated Cdh5-Cre^ERT2^(PAC)/ZsGreen mice and examined whether these ZsGreen⁻ hematopoietic progenitors can become fluorescent after tamoxifen administration (Figure 4A). As a positive control, we also transplanted lethally irradiated WT C57Bl6 mice (n=2) with an LSK population enriched for ZsGreen⁺ cells (45.9% purity, with the remaining 54.1% comprising ZsGreen^-^ LSK cells (Figure 4A and Figure S3F). Four weeks post-transplant, we administered tamoxifen and monitored the peripheral blood for the presence of ZsGreen^+^ hematopoietic cells over six months.

**Figure 4.**
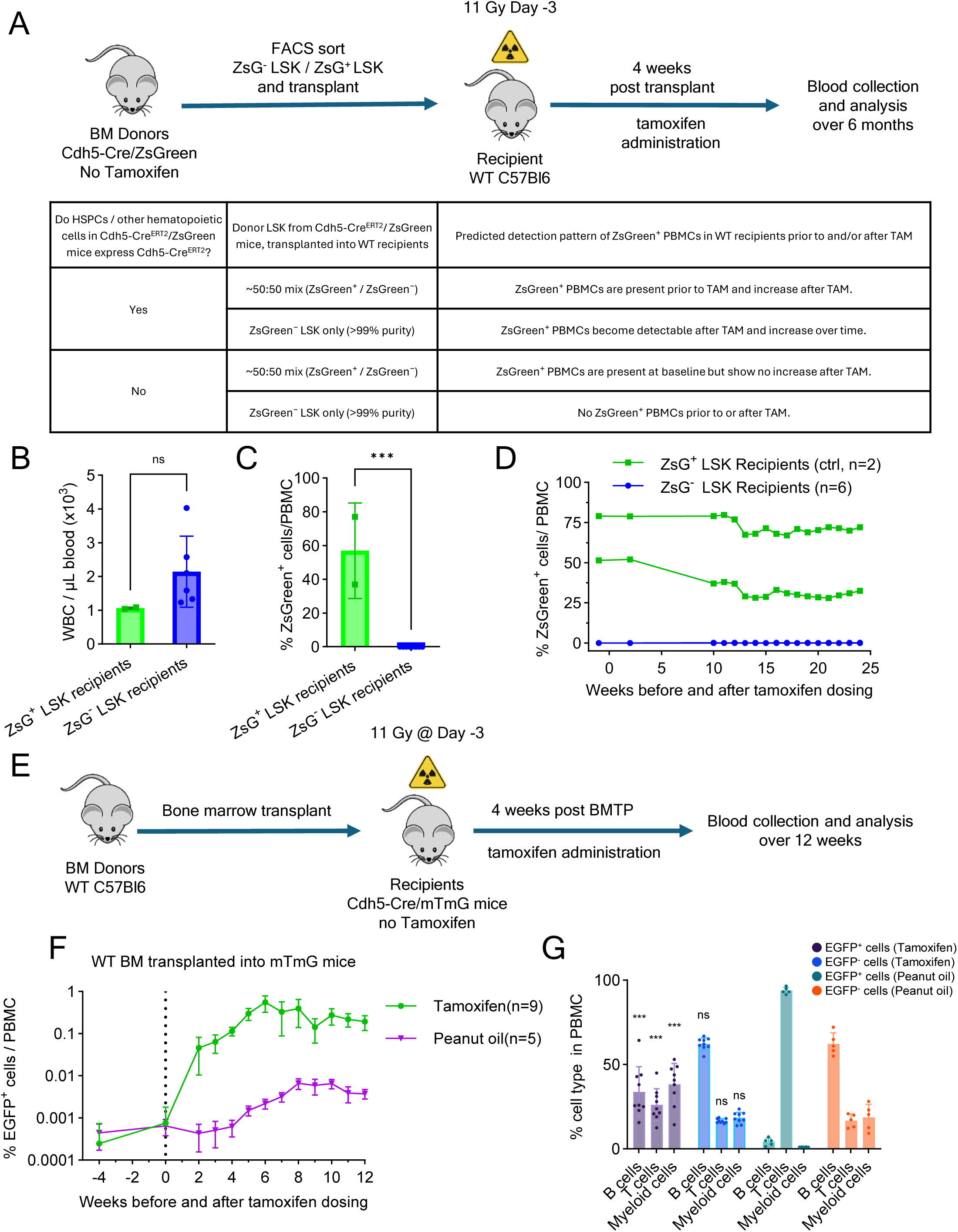
Independence of adult EHT from preexisting HSPC (A) Transplantation experiment: donor LSK sorting, recipient irradiation, transplantation, tamoxifen treatment, and analysis (top). Tabular representation of possible outcomes of the experiment designed to address the question “Do HSPCs/other hematopoietic cells in Cdh5Cre^ERT2^/ZsGreen mice express Cdh5-Cre^ERT2^?” (bottom). (B-D) Blood WBC counts (B), percent ZsGreen⁺ PBMC (C), and time course of ZsGreen⁺ PBMC detection (D) in transplant recipients of ZsGreen⁻ LSK (5×10^4^ or 2.5×10^4^ cells/mouse; n=3/group) and ZsGreen-enriched LSKs (2.8×10^3^ cells/mouse; n=2). Results in B and C are from week 24 post-tamoxifen. (E) Experiment: WT BM transplantation (BMTP) into lethally irradiated Cdh5-Cre/mTmG mice (n=9). Four weeks later, tamoxifen was administered; blood was monitored for 16 weeks. (F and G) EGFP⁺ PBMC detection before and after tamoxifen or peanut oil administration (F) and cell type distribution of EGFP⁺ and EGFP⁻ PBMCs at week 12 post-tamoxifen or peanut oil (G) in Cdh5-Cre/mTmG recipients (n=9) of WT BM (5×10^6^ cells). Statistical significance reflects comparisons between EGFP^+^ and EGFP^-^ cells in the tamoxifen vs peanut oil groups. Dots represent individual mice. Data are shown as mean ± SD. ***p < 0.001, ns, not significant by Student’s *t* test.

Expectedly, all LSK recipients (ZsGreen^-^ LSKs or LSK enriched with ZsGreen^+^ cells) showed successful hematopoietic reconstitution as evidenced by normal blood WBC counts at 10 weeks post-transplant (Figure 4B). Importantly, the 6 mice transplanted with ZsGreen⁻ LSKs did not produce ZsGreen⁺ PBMCs post-tamoxifen administration, indicating that the transplanted LSKs and their progeny did not express tamoxifen inducible Cdh5-Cre^ERT2^ recombinase activity (Figure 4C and 4D). Instead, the 2 mice transplanted with ZsGreen⁺-enriched LSKs displayed a similar percent of ZsGreen⁺ PBMCs prior to and after tamoxifen administration (Figure 4D).

Collectively, these results demonstrate that LSK progenitors in adult BM lack of tamoxifen-inducible Cdh5 expression and do not contribute to tamoxifen-induced adult EHT in our Cdh5-reporter mice. Rather, these results strongly support the conclusion that Cdh5^+^CD45^-^ ECs are a source of hematopoietic cells in the adult mouse.

In additional experiments, we took advantage of the relative insensitivity of BM ECs subsets to irradiation relative to BM hematopoietic cells^50^ to examine the possibility that EC surviving after lethal irradiation may be hemogenic. As a lethal dose of irradiation effectively eliminates HSPCs and requires hematopoietic reconstitution for survival, we transplanted WT (untracked) BM cells into lethally irradiated Cdh5-Cre/mTmG mice and treated the mice with tamoxifen 4 weeks after transplantation (Figure 4E). Prior to tamoxifen administration, >99% of PBMCs were not fluorescent, indicating that these cells derived from the transplanted WT BM rather than host-derived (Figure 4F). After tamoxifen treatment, a progressive increase in EGFP⁺ PBMCs was observed, reaching ∼0.55% by six weeks, which included myeloid cells and B and T lymphocytes (Figure 4F and 4G). Although we cannot exclude the possibility that rare EGFP^+^ hematopoietic progenitor (tracked tamoxifen-dependently or independently) may have survived the irradiation, the presence of EGFP⁺ hematopoietic cells in the circulation of lethally irradiated Cdh5-Cre/mTmG mice suggests their derivation from radioresistant Cdh5^+^ ECs rather than from radiosensitive hematopoietic progenitors. These results further support the view that adult ECs possess hemogenic potential and can produce hematopoietic cells in vivo.

### Single cell tracking confirms the presence hemogenic ECs in adult BM

To directly trace hematopoietic cell progeny arising from individual adult ECs, we exploited the *PolyloxExpress* sc genetic barcoding system. We generated Cdh5-Cre^ERT2^/ZsGreen/*PolyloxExpress* mice, in which both the ZsGreen and Polylox transgenes are inserted in the Rosa26 locus, such that individual ZsGreen^+^ cells contain a single Polylox barcode^33,51,52^. To evaluate EC-derived hematopoietic cell output, we harvested BM from tamoxifen-treated Cdh5-Cre^ERT2^/ZsGreen/*PolyloxExpress* mice (n=3), sorted the ZsGreen^+^VE-Cadherin^+^Endomucin^+^ (purity >95%) and the ZsGreen^+^VE-Cadherin^-^Endomucin^-^CD45^+^ hematopoietic cells (purity >98%), mixed these populations (1:1 ratio; total 147,446 cells) and processed for 10x Illumina Sequencing and PacBio sequencing (detailed in Methods). We recovered sc transcriptome from 93,553 cells; of these, 4,069 cells had a barcode (Figure 5A, Figure S5A-B, detailed in Methods).

**Figure 5.**
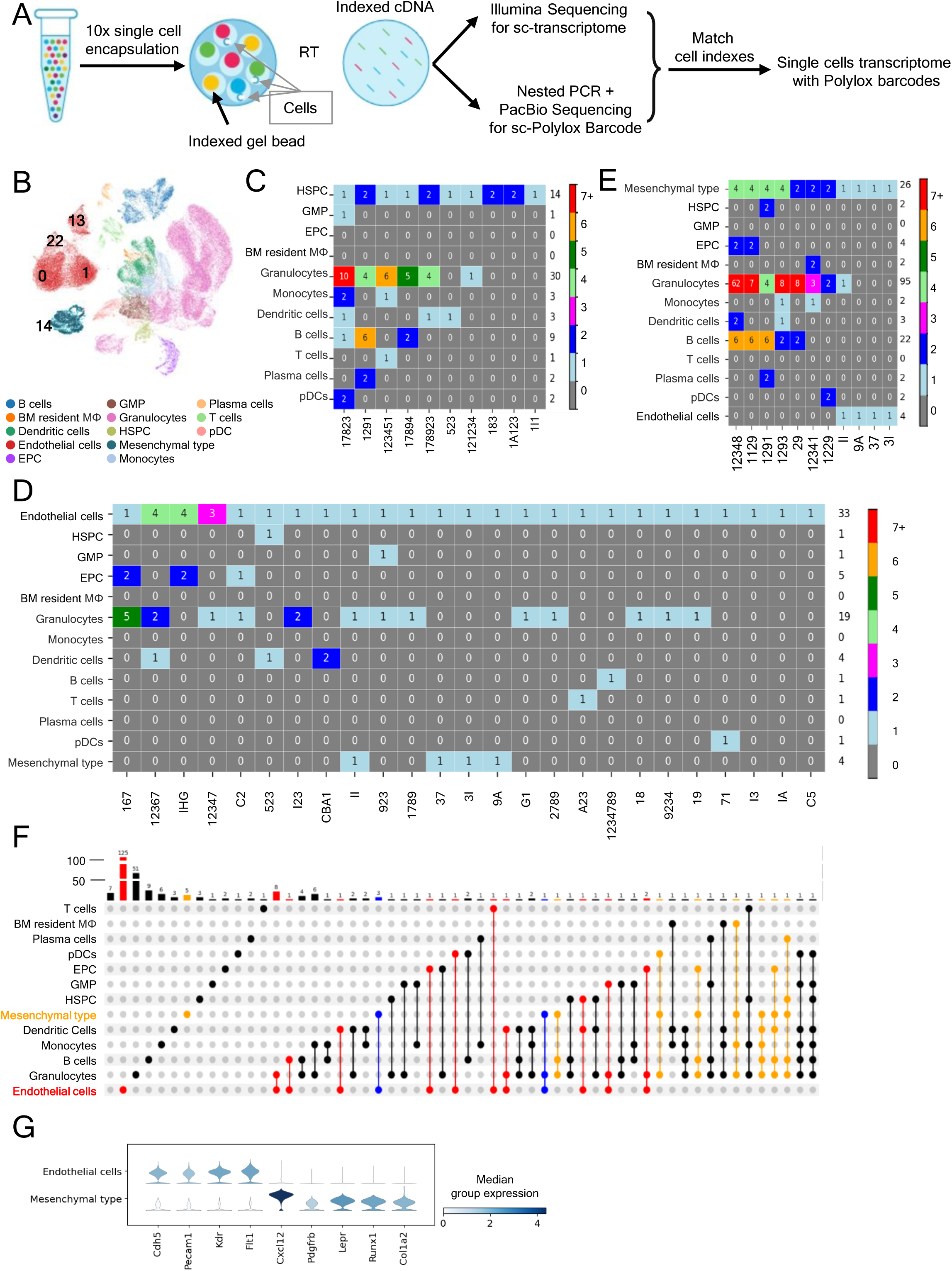
Polylox sc lineage tracing links adult BM ECs to hematopoietic progenitors and mature blood cell progeny (A) Schematic of Polylox barcode and transcriptome profiling. FACS-enriched ECs (ZsGreen⁺VE-Cadherin^+^Endomucin^+^) and EC-depleted (ZsGreen^-^VE-Cadherin^-^Endomucin^-^) BM cells from tamoxifen-treated Cdh5-Cre^ERT2^/ZsGreen/PolyloxExpress mice (n=3, 10-week-old at the time of tamoxifen treatment) were mixed (1:1), and encapsulated (147,446 cells loaded; 93,553 processed). Indexed cDNA was used for scRNA-seq and barcode detection by PacBio sequencing after nested PCR enrichment; barcode-transcriptome integration was accomplished via shared cell indices. (B) UMAP clustering and cell type annotation. Clusters 0, 1, 13, and 22 comprise ECs; cluster 14 comprises Mesenchymal-type cells. (C – E) Heatmaps showing “true” Polylox barcodes (pGen < 1×10⁻^6^) linking HSPCs to hematopoietic cells (C), ECs to hematopoietic and other cells (D), and Mesenchymal-type cells to other cells (E). The numbers within the colored boxes identify cell number; the labels at the bottom of each column denote the barcode shared by all cells in that column; the number on the right side the heatmaps reflects the total number of cells in each row. (F) UpSet plot showing cells (identified by colored dots) sharing the same “true” barcode (identified by lines connecting the colored dots); bar graph at the top of the plot reflects (height and number on each bar) the number of “true” barcodes. Colors of dots: EC (red), Mesenchymal-type (orange), ECs connecting with Mesenchymal-type cells (blue), cells other than ECs and Mesenchymal-type cells (black). (G) Violin plots showing selected gene expression profile in Mesenchymal-type cells (cluster 14) and ECs (clusters 0, 1, 13, 22 combined).

Unsupervised clustering of sc transcriptome data revealed 34 clusters, 31 of which remained after doublet removal (Figure 5B; Figure S4A-S4D). Cell clusters 0, 1, 22, and 13 were annotated as “Endothelial cells” based on expression of the classical EC markers *Cdh5*, *Pecam1*, *Kdr* and *Flt1*, and absence of *Ptprc*/CD45, *Runx1*, and the mesenchymal cell markers *Cxcl12*, *Lepr*, *Pdgfrb*, and *Col1a2* expression (Figure 5B; Figure S4E). Cell cluster 14 was annotated as “Mesenchymal type” based on co-expression of *Cxcl12*, *Lepr*, *Pdgfrb*, *Col1a2*, but expressed *Runx1* and the EC markers *Cdh5* and *Pecam1* (Figure 5B; Figure S4E). The remaining cell clusters included the hematopoietic progenitors and mature blood cells of various lineages (Figure 5B; Figure S4E). Cre^ERT2^ transcripts were detected exclusively in Cdh5-expressing endothelial cell populations and were absent from Ptprc/CD45-expressing hematopoietic cells (Figure S4E). Cre^ERT2^ expression levels in ECs were low but closely tracked with the expression patterns of canonical endothelial markers (*Cdh5*, *Pecam1*, *Emcn*, and *Eng*).

Barcode analysis revealed robust *Polylox* barcode diversity among cell populations, including 274 “true” barcodes, defined as barcodes with low generation probability (Figure S5A, *P* _gen_ <1×10^-6^)^51^ consistent with rare, unique recombination events (Figure S5A - S5C). Expectedly, “true” barcodes linked HSPC to downstream progenitors and mature hematopoietic cells, validating the system (Figure 5C).

Notably, 169 of 828 ECs (from “endothelial” cell clusters 0, 1, 22, and 13) were marked with “true” barcodes. The results detected 19 links between EC and hematopoietic cells based on their shared “true” barcode (Figure 5D). These hematopoietic cells linked to ECs by shared “true” barcodes included HSPC, EPC, GMP, and mature blood cells, encompassing granulocytes, monocytes, dendritic cells, B and T lymphocytes, plasma cells, and pDCs (Figure 5D and E). These results provide direct evidence, at sc resolution, that adult mouse BM ECs can generate hematopoietic progenitors and mature blood cells.

Additionally, 26 “Mesenchymal type” cells (from cluster 14; co-expressing mesenchymal cell markers, *Runx1*, and the EC markers *Cdh5* and *Pecam1*) were also marked by 11 “true” barcodes, 8 of which were shared with hematopoietic cell progenitors (HSPC and Erythroid) and mature blood cells (Figure 5E and F). Also, 4 “true” barcodes linked “Mesenchymal type” cells to ECs (from clusters 0, 1, 22 and 13), suggesting either a shared precursor or derivation from each other. Although it cannot be excluded that barcoding missed identification of mesenchymal cell links to other ECs or cells, these results raise the possibility that certain BM “Mesenchymal type” cells may produce hematopoietic cell progeny. Despite similarities of sc tracing results linking ECs and “Mesenchymal type” cells to hematopoietic cells, these two cell populations display a distinctive transcriptome profile (Figure 5G; Figure S4E).

Additional analysis of all tracked cells showed that several ECs and to a lower extent “Mesenchymal type cells” shared “true” barcodes (Figure S5D-F), indicating clonal expansion. Transcriptome-based sc cell cycle analysis, confirmed the presence of ECs and “Mesenchymal type cells” in the S and G2/M phases, albeit to a much lower degree than HSPC (Figure S5C).

Together, these results demonstrate at a sc level that adult BM ECs can generate hematopoietic cell progeny of HSPC and mature blood cells. The results further raise the possibility that “Mesenchymal type” cells, marked by a hybrid endothelial and stromal phenotype, may represent an additional source of hemogenic activity in adult BM.

### Single cell transcriptome identifies a *Cdh5*^+^*Col1a2*^+^ *Runx1*^+^cell population in the adult BM

To further characterize adult hemogenic cell populations, we analyzed publicly available scRNAseq datasets comprising BM cells from adult mice (1-16 months of age)^53–59^ and embryonic caudal artery ECs (9.5–11.5 days post coitum)^24^. After quality control and dimensional reduction, the remaining 434,810 cells clustered into 71 distinct populations (Figure 6A-D). Among the *Cdh5*-expressing endothelial clusters, two clusters, cluster 8 composed predominantly of embryonic cells (98%) and cluster 50 composed largely of adult BM-derived cells (95.5%) (Figure 6E), were notable in comprising cells co-expressing *Cdh5* and *Runx1*, a transcription factor that marks the hemogenic EC identity during development^60^ (Figure 6B).

**Figure 6.**
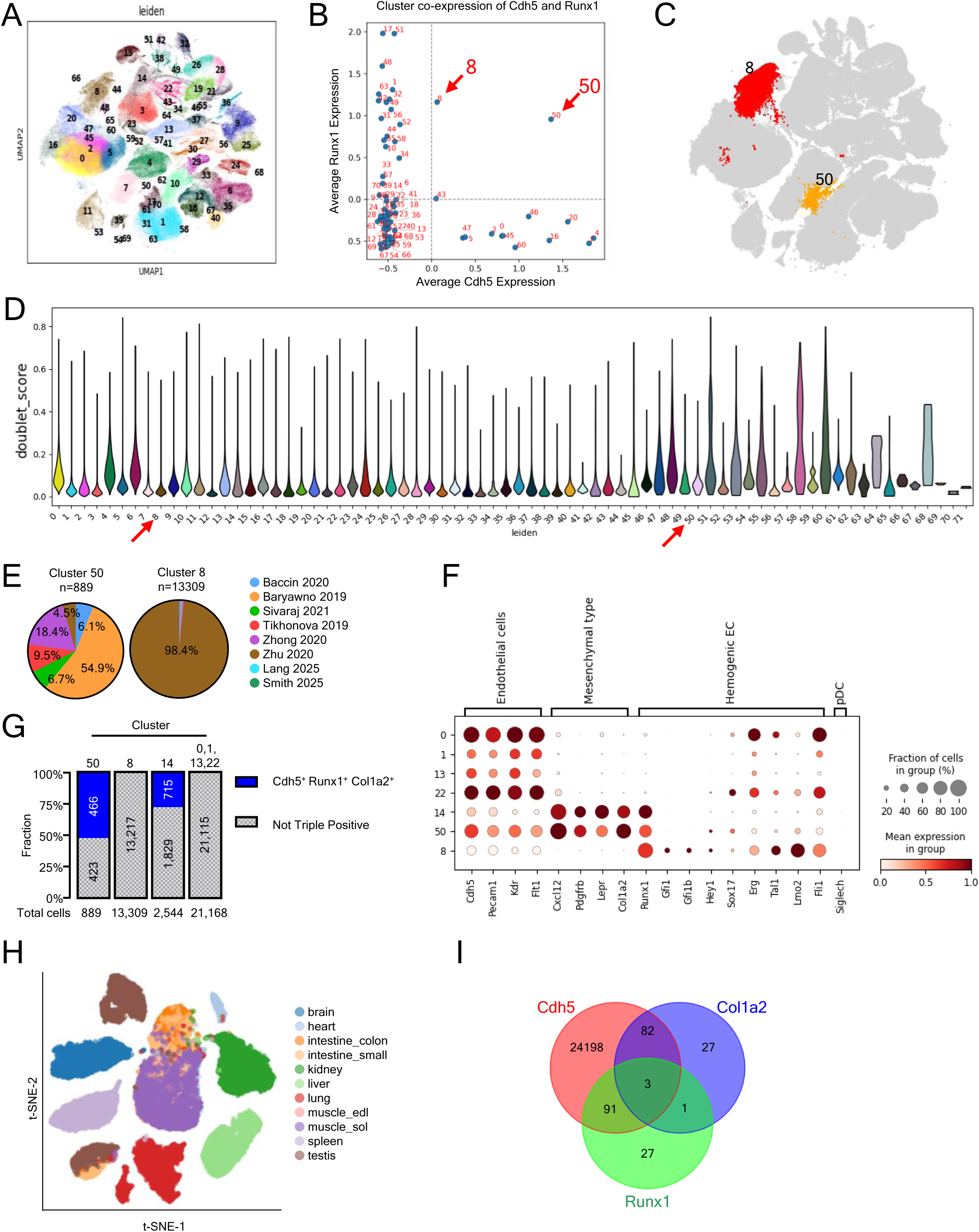
Sc transcriptomic analysis of prospective hemogenic ECs (A) UMAP clustering of 434,810 cells from eight public scRNA-seq datasets. (B) Dot plot showing relative Cdh5 and Runx1 co-expression across clusters; clusters 8 and 50 co-express both genes. (C) UMAP highlighting clusters 8 and 50; all other clusters shown in grey. (D) Violin plots of doublet scores across Leiden clusters. Clusters 50 and 8 show no evidence of doublet enrichment. (E) Datasets proportional contribution to clusters 50 and 8; each dataset is color-coded. (F) Dot plot showing expression of selected marker genes in clusters 50 and 8 (from the public sc RNA-seq datasets listed in Figure 7D) and from clusters 0, 1, 13, 22 and 14 (from Polylox scRNA-seq; Figure 5B). Results reflect mean expression and fraction of cells in group. (G) Cdh5, Runx1 and Col1a2 co-expression in the indicated clusters as a fraction of cells in the cluster. (H and I) t-SNE plot of ECs from 11 murine tissues (G) and Venn diagram (H) showing rare co-expression of Cdh5, Runx1, and Col1a2 in these tissues.

We jointly analyzed the transcriptome from the publicly available embryonic cluster 8 and adult cluster 50 (Figure 6 A-C), and from our sc Polylox dataset, including adult clusters 14 (Mesenchymal type) and adult EC clusters 0, 1, 13 and 22 (Figure 5B). All populations expressed canonical EC markers (*Cdh5*, *Pecam1*, *Kdr*, *Flt1*), though at different levels, but expression of *Runx1* was mostly confined to clusters 8, 50 and Polylox 14 (Figure 6F).

Interestingly, adult clusters 50 and Polylox 14 co-expressed the mesenchymal cell-associated genes *Col1a2*, *Lepr*, *Cxcl12*, and *Pdgfrb*, distinguishing these two clusters from the embryonic cluster 8 and adult Polylox clusters 0, 1, 13 and 22 (Figure 6F). Additionally, embryonic cluster 8 exhibited higher expression of key EHT-related transcription factors (FLI1, LMO2, TAL1, ERG)^61–64^, some of which were variably expressed by the Polylox clusters 0,1,13 and 22 (Figure 6F).

To explore whether cells expressing *Cdh5*, *Col1a2* and *Runx1* are unique to BM (Figure 6G), we looked at other adult mouse tissues. Analysis of public scRNAseq datasets from 11 adult mouse tissues^65^ showed that *Cdh5*, *Runx1*, and *Col1a2* expressing ECs are largely restricted to the BM, with only 3 such cells detected out of ∼32,000 EC from other tissues (Figure 6H and I)^65^. These observations, together with the Polylox sc tracing experiments, indicate that adult mouse BM harbors a cell population with mixed endothelial and mesenchymal phenotype marked by *Runx1*, *Col1a2* and *Cdh5* expression and raises the possibility that this population has hemogenic potential.

### *Col1a2* and *Runx1* expression in BM ECs

To evaluate a potential contribution of *Col1a2* expression to EHT in adult BM, we generated the mouse lines Col1a2-Cre^ERT2^/mTmG and Col1a2-Cre^ERT2^/ZsGreen to track cells expressing the *Col1a2* gene (Figure S6A). Four weeks after tamoxifen administration, the BM of Col1a2-Cre^ERT2^/ZsGreen mice contained abundant ZsGreen^+^ cells (Figure S6B). By flow cytometry, a subset of these BM Col1a2-tracked ZsGreen^+^ cells were VE-Cadherin^+^CD45^-^ and RUNX1^+^ consistent with an endothelial identity and perhaps hemogenic potential (Figure S6C).

Noteworthy, a similar subset of VE-Cadherin^+^CD45^-^RUNX1^+^ cells was detected in BM of adult WT C57Bl/6 mice (Figure S6D) and in BM of Cdh5-Cre^ERT2^(PAC)/ZsGreen mice after tamoxifen treatment (Figure S6E).

To evaluate hemogenic potential, we looked for Col1a2-tracked CD45^+^ hematopoietic cells in Col1a2-Cre^ERT2^/mTmG and Col1a2-Cre^ERT2^/ZsGreen mouse lines after tamoxifen treatment. In both these mouse lines, we detected CD45^+^ hematopoietic cells tracked by EGFP or ZsGreen fluorescence in BM and blood, which were rare but significantly more abundant than in control mice not treated with tamoxifen (Figure 7A and B; Figure S6F-H). These results indicated that a proportion of the Col1a2-tracked cells in the adult mouse are hemogenic.

**Figure 7.**
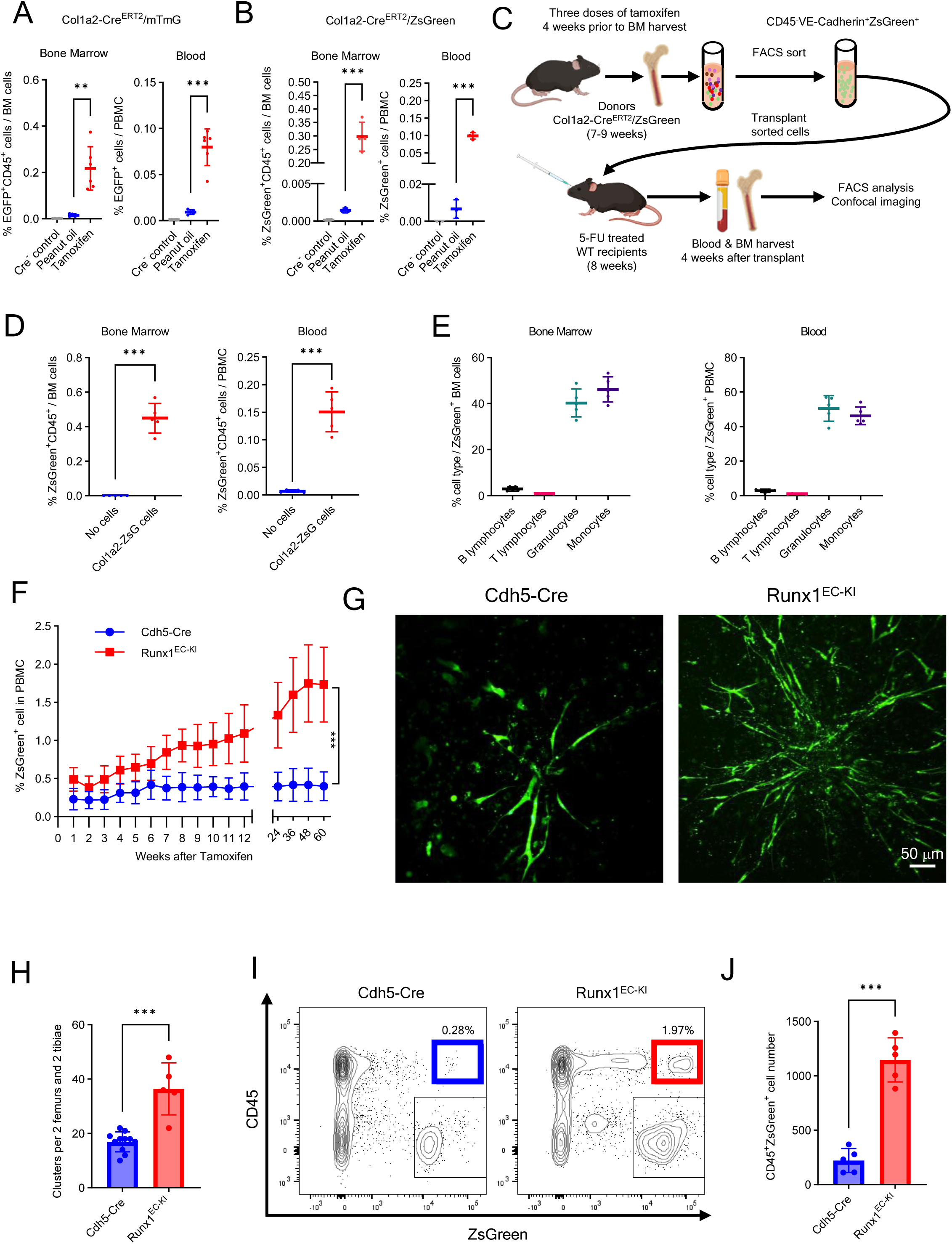
Contribution of *Col1a2* and *Runx1* expression to ECs hemogenic activity (A and B) Percent EGFP^+^CD45^+^ cells in BM and blood of tamoxifen-treated (n=6) or oil-treated (n=5) Col1a2-Cre^ERT2^/mTmG mice (A) and tamoxifen-treated (n=4) or oil-treated (n=3) Col1a2-Cre^ERT2^/ZsGreen mice (B). Cre-control mice (n=5 in A, and n=2 in B). (C) Transplant experiment: sorted VE-Cadherin⁺CD45⁻ZsGreen⁺/Col1a2⁺ cells from tamoxifen-treated Col1a2-Cre^ERT2^/ZsGreen mice are transplanted into 5-FU-conditioned WT recipients. (D and E) Detection (D) and characterization (E) of ZsGreen⁺CD45⁺ cells in BM and blood of WT 5-FU-conditioned mice (n=5), 4 weeks post-transplant of VE-Cadherin⁺CD45⁻ZsGreen⁺/Col1a2⁺ cells. Control FU-conditioned WT mice (n=4) received no cell transplant (D). (F) Time course of ZsGreen⁺ PBMC detection in control (Cdh5-Cre⁺/ZsGreen⁺) and Runx1^EC-KI^ (Cdh5-Cre⁺/ZsGreen⁺/Runx1-KI) mice (n=10 per group). (G and H) Representative images (G) and quantification (H) of ZsGreen⁺ cells from OP9 cell-supported cultures of BM cells from tamoxifen-treated Cdh5-Cre⁺/ZsGreen⁺ (n=11) and Runx1^EC-KI^ mice (n=5). (I and J) Representative flow cytometry plots (I) and quantification (J) of CD45⁺ZsGreen⁺ cells from OP9 cell-supported BM cell cultures (n=5/group). Dots represent individual mice. Data are shown as mean ± SD. **p<0.01, ***p < 0.001 by Student’s t test.

To further evaluate this possibility, we first sorted VE-Cadherin^+^CD45^-^ ZsGreen (Col1a2)^+^ cells from the BM of tamoxifen-induced adult Col1a2-Cre^ERT2^/ZsGreen mice (Figure S7A), examined expression of selected genes, and used these cells in transplant experiments. The sorted VE-Cadherin^+^CD45^-^ ZsGreen (Col1a2)^+^ cells expressed *Cdh5*, *Col1a2*, *Cxcl12*, and *Runx1* mRNAs distinctively from other BM cell population (Figure S7B), but resembled the BM hemogenic population annotated as “mesenchymal-type” (cluster 14) identified by Polylox s.c sequencing (Figure 5G; Figure S4E) and subsets of adult BM Cdh5^+^ cells identified in public adult datasets (Figure 6E and F).

We transplanted the BM VE-Cadherin^+^CD45^-^ ZsGreen (Col1a2)^+^ cells (1×10^4^ cells/mouse) into 5-FU-conditioned adult WT C57Bl/6 recipients and looked for tracked CD45^+^ cells in the BM and blood (Figure 7C). Four weeks after transplantation, BM and peripheral blood of transplant recipients contained ZsGreen^+^CD45^+^ cells (Figure 7D; Figure S7C and D), indicating that the transplanted ZsGreen^+^ (Col1a2)-tracked CD45^-^ ECs had produced CD45^+^ hematopoietic cell progeny. These ZsGreen^+^CD45^+^ cells in the recipient mice comprised mainly granulocytes and monocytes, and few B and T lymphocytes (Figure 7E). These results indicate that Col1a2-tracked ECs, like Cdh5(VE-Cadherin)-tracked ECs, can give rise to hematopoietic progeny in vivo, but display a more restricted multilineage potential compared to ECs from Cdh5-Cre^ERT2^(PAC)/ZsGreen mice.

In additional experiments, we evaluated the role of *Runx1* expression in adult BM hemogenic EC since our analyses identified *Runx1* as a putative marker of adult EHT (Figure 6E and F; Figure S4E). Therefore, we generated a Runx1^EC-KI^ mouse line (*Cdh5-Cre^ERT2^/ZsGreen/Runx1-Knock-In*), in which Cre-mediated excision of a floxed STOP codon enables co-expression of ZsGreen and *Runx1* in ECs upon tamoxifen induction (Figure S7E)^66^. In these Runx1^EC-KI^ mice, tamoxifen treatment increased significantly the frequency of ZsGreen⁺ cells in PBMC over 60 weeks compared to control tamoxifen-treated (Cdh5-Cre^ERT2^/ZsGreen) mice (Figure 7F).

Since these circulating ZsGreen-tracked cells presumably represent hematopoietic cell progeny of ECs co-expressing ZsGreen and *Runx1*, we tested the hemogenic potential of these tamoxifen-induced BM EC ex vivo. Using the culture system that successfully supported ex vivo hematopoiesis in Cdh5-tracked BM (Figure 2A-H), we now compared the hemogenic potential of BM cells from tamoxifen pretreated Runx1^EC-KI^ mice to that of BM cells from tamoxifen pretreated Cdh5-Cre^ERT2^/ZsGreen) mice. We detected a significantly greater number of ZsGreen^+^ clusters in cultures from Runx1^EC-KI^ BM cells compared to control BM cells (Figure 7G and H), and flow cytometry showed that more CD45⁺ZsGreen⁺ hematopoietic cells were produced in cultures of Runx1^EC-KI^ BM cells compared to BM from controls (Figure 7I and J). Although we cannot exclude the possibility that RUNX1 promotes EC proliferation in culture, these results support a role of RUNX1 in promoting adult EHT.

## DISCUSSION

Our results provide evidence for the presence of ECs in the adult mouse BM with hemogenic potential. Previously, hemogenic ECs were detected during embryonic development or perinatally but not thereafter^1-9,67^. The current findings argue that EHT is not limited to the prenatal or perinatal development but is present up to 28 weeks after birth, decreasing thereafter. This conclusion is supported by Cdh5-based bulk and Polylox sc lineage tracking, culture of hemogenic ECs, transplant analyses and initial characterization of EC-derived hematopoietic cell progeny, which link features of adult EHT to embryonic EHT^25–28,68^.

Previous experiments found that EHT, present in the late fetus/neonatal BM, disappears shortly after birth^67^. This contrasts with the current experiments showing persistence of adult EHT well beyond 20 weeks of age. The divergent results likely stem from functional differences of the Cdh5-based tracking systems. In the previous experiments, attempts to activate Cdh5-fluorescence in BM ECs by injecting tamoxifen at different time-points after birth were unsuccessful starting 20 days after birth. Expectedly, the absence of tamoxifen-induced fluorescence in BM ECs was associated with the absence of traced hematopoietic cell output from these cells. In the current experiments and those of others^36^, tamoxifen administration after 10, 20, and 30 weeks of age consistently induces fluorescence in most BM ECs. We conclude that the absence of adult EHT had not been firmly established.

Adult hemogenic ECs identified here express *Cdh5*, *Pecam1*, *Kdr*, and *Flt1*, resembling other BM EC populations and embryonic hemogenic ECs^69^. However, the current studies identify yet another, small hemogenic cell population in the adult BM, distinctive in co-expressing typical EC markers, *Cdh5* and *Pecam1*, and the mesenchymal-type markers *Lepr*, *Col1a2* and *Cxcl12*. These two hemogenic populations are clonally linked, but their relationship is currently unclear.

ECs derive from two sources during development: the splanchnic mesoderm that gives rise to the primitive aorta where ECs located on the aortic floor are hemogenic^68^, and the somites from the paraxial mesoderm, that produce ECs contributing the endothelial vascular network of the trunk and limb^70,71^. Somite-derived ECs are not hemogenic in situ, but they can transiently acquire a hemogenic potential when variously induced^72^ and may also include a cell subset with hemogenic potential^73^. They can also express *Runx1* and trigger aortic hematopoiesis from hemogenic ECs in zebrafish^74^ and in the chick^68^. Besides generating hemogenic ECs, blood and other tissues, the mesoderm in conjunction with the neural crest, gives rise to mesenchymal stem cells (MSCs) and perhaps a precursor of both MSCs and ECs^75–77^. However, MSCs do not produce blood cells, although they can differentiate into many other tissues^78^, and their potential for endothelial differentiation remains controversial^79–82^. By contrast, endothelial to mesenchymal transition (EndMT) is a well-established process in development and disease states^83^.

It was suggested that the transient wave of EHT occurring in the BM of the late fetus and perinatally may serve to mitigate the slow HSC colonization of BM from the fetal liver or perhaps prepare the BM niches to accommodate the incoming HSC from the liver, but a function has not yet been firmly established^67^. Similarly, the function of adult BM EHT identified here is currently unclear. We hypothesize that adult EHT plays a functional role under conditions of hematopoietic stress or disease rather than under steady-state conditions, but this will need future investigation. Interestingly, previous studies found that subsets of ECs can regenerate after irradiation that has eliminated HSC^45^, and we traced the emergence of some hematopoietic cells from putative ECs that persisted after lethal mouse irradiation. We also found that 5-FU treatment was required for the successful engraftment and function of adult hemogenic ECs, suggesting that this population may require inductive signals from a BM that is recovering from an insult^45,84^. The identification of such inductive factors may enable effective propagation of BM hemogenic ECs ex vivo and motivate a search for hemogenic ECs in human BM.

Altogether, our results point to a previously unrecognized capability of ECs in the adult mouse BM to generate blood cells. These results suggest that hematopoiesis in the adult mouse may arise through the contribution of cells and processes beyond the HSCs generated through aortic EHT during development and BM EHT perinatally.

### Limitations of the study

A limitation of our study is that the function of adult EHT is currently unknown. Under steady-state conditions, the hematopoietic cell output from adult EHT is very low in comparison to that of embryonic EHT, which effectively sustains adult hematopoiesis, and our experiments detected no functional differences between hematopoietic cells from embryonic and adult EHT. To address this limitation, we will examine whether adult EHT is a greater contributor to adult hematopoiesis when stress is imposed on the adult hematopoietic system. In addition, we have not fully characterized the hematopoietic cell population(s) arising from adult EHT or studied the time-course kinetics of hematopoietic cell emergence. A key issue is whether long-term repopulating HSPCs are produced by adult EHT. This will require serial transplantation experiments, including CD45.1/CD45.2 competitive repopulation assays, and kinetic analysis of their emergence from hemogenic ECs. Finally, we could not extend the in vivo single-cell PolyloxExpress genetic lineage tracing to the ex vivo single-cell genetic lineage tracing due to technical limitations of the PolyloxExpress system.

## RESOURCE AVAILABILITY

### Lead contact

Further information and requests for resources and reagents should be directed to and will be fulfilled by the lead contact, Giovanna Tosato.

### Materials availability

This study did not generate new unique reagents.

## Data and code availability

The results of scRNAseq and PacBio SmrtSeq of Polylox barcodes are deposited in NCBI SRA (PRJNA1079369, public at time of publication). A custom script was used to retrieve cell indexes from the PolyloxExpress amplicons (https://github.com/CCRSF-IFX/SF_Polylox-BC). Bash, R, and Python codes are available from the corresponding author upon reasonable request.

## ACKNOWLEDGEMENTS

This project is supported by the Intramural Program of CCR, NCI, NIH. The findings are those of the authors and do not necessarily represent the official views of the NIH or the Department of Health and Human Services. This work used the computational resources of the NIH High Performance Computing (HPC) Biowulf cluster (http://hpc.nih.gov) and Frederick Research Computing Environment (FRCE). Flow cytometry and cell sorting were performed at the CCR Flow Cytometry Core Facility; microscopy analyses at the CCR Microscopy Core Facility in Building 37 of the NCI, NIH. We thank Drs. Ralf Adams and Manfred Boehm for mouse line *Cdh5-Cre^ERT2^(PAC)*; Drs. Yoshiaki Kubota and Yosuke Mukoyama for mouse line *Cdh5-Cre^ERT2^(BAC)*; Dr. Hans-Reimer Rodewald for the *PolyloxExpress* mouse line. We thank Dr. S. Banerjee, Dr. S. Siddiqui and Ms. K. M. Wolcott for flow cytometry support; Dr. M. Kruhlak and Mr. L. Lim for confocal microscopy support and Mr. A. Abdelmaksoud for bioinformatics assistance. We thank Drs. D. Lowy, R. Yarchoan, M. DiPrima and H. Ohnuki for thoughtfully commenting on aspects of this work.

## AUTHOR CONTRIBUTIONS

Concept and design: J-X.F. and G.T. Acquisition, analysis, or interpretation of data: J-X.F., G.T., C.L., F.L., N.T., A.B., L.L, M-T.Y., D.W., J.C., and Y.Z. G. Drafting of the manuscript: J-X.F., G.T. Critical revision of the manuscript for important intellectual content: J-X.F., G.T., A.B. and N.T.

## SUPPLEMENTAL INFORMATION

Document S1. Figures S1–S7

## METHODS

### RESOURCES TABLE

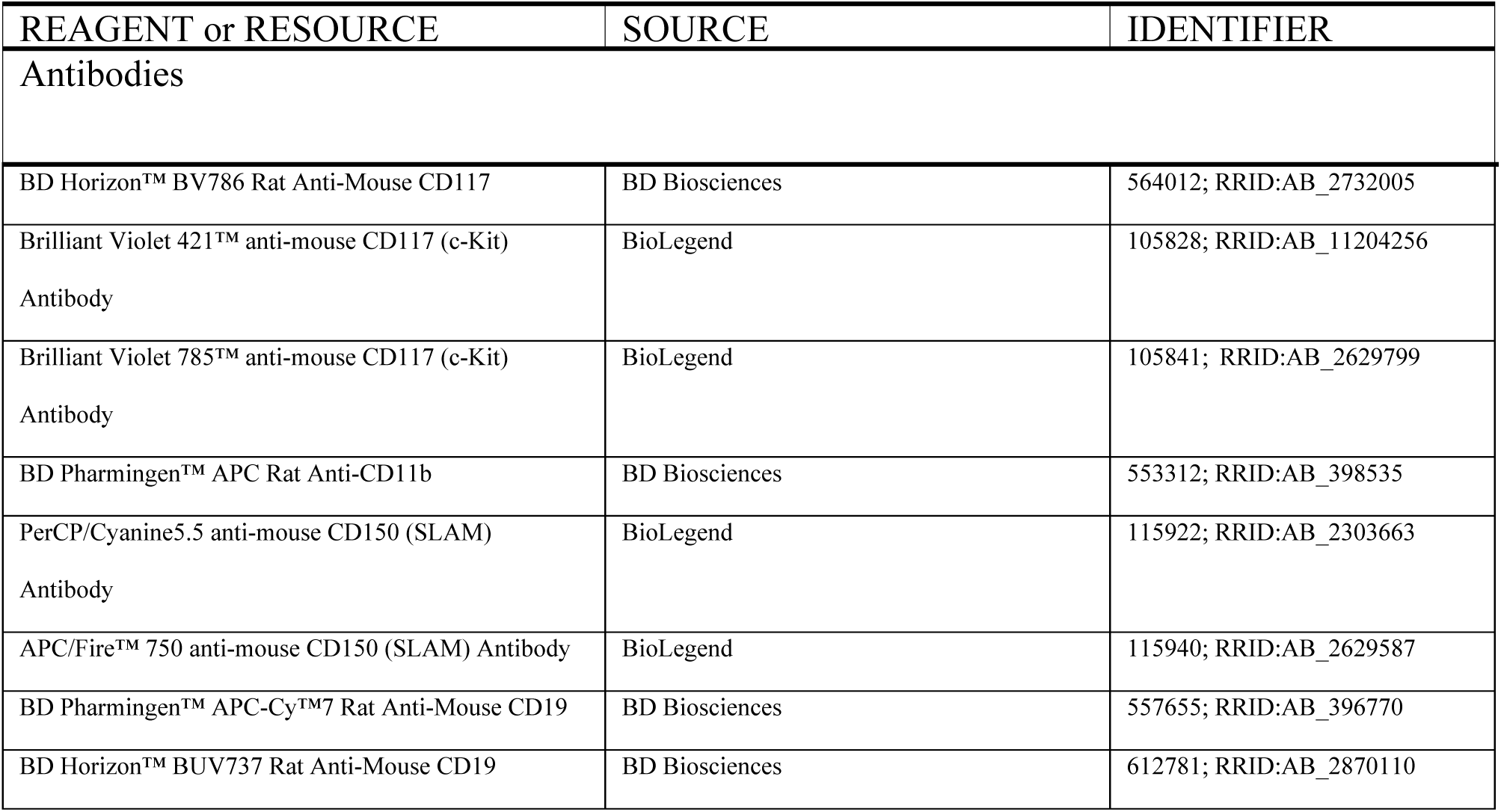
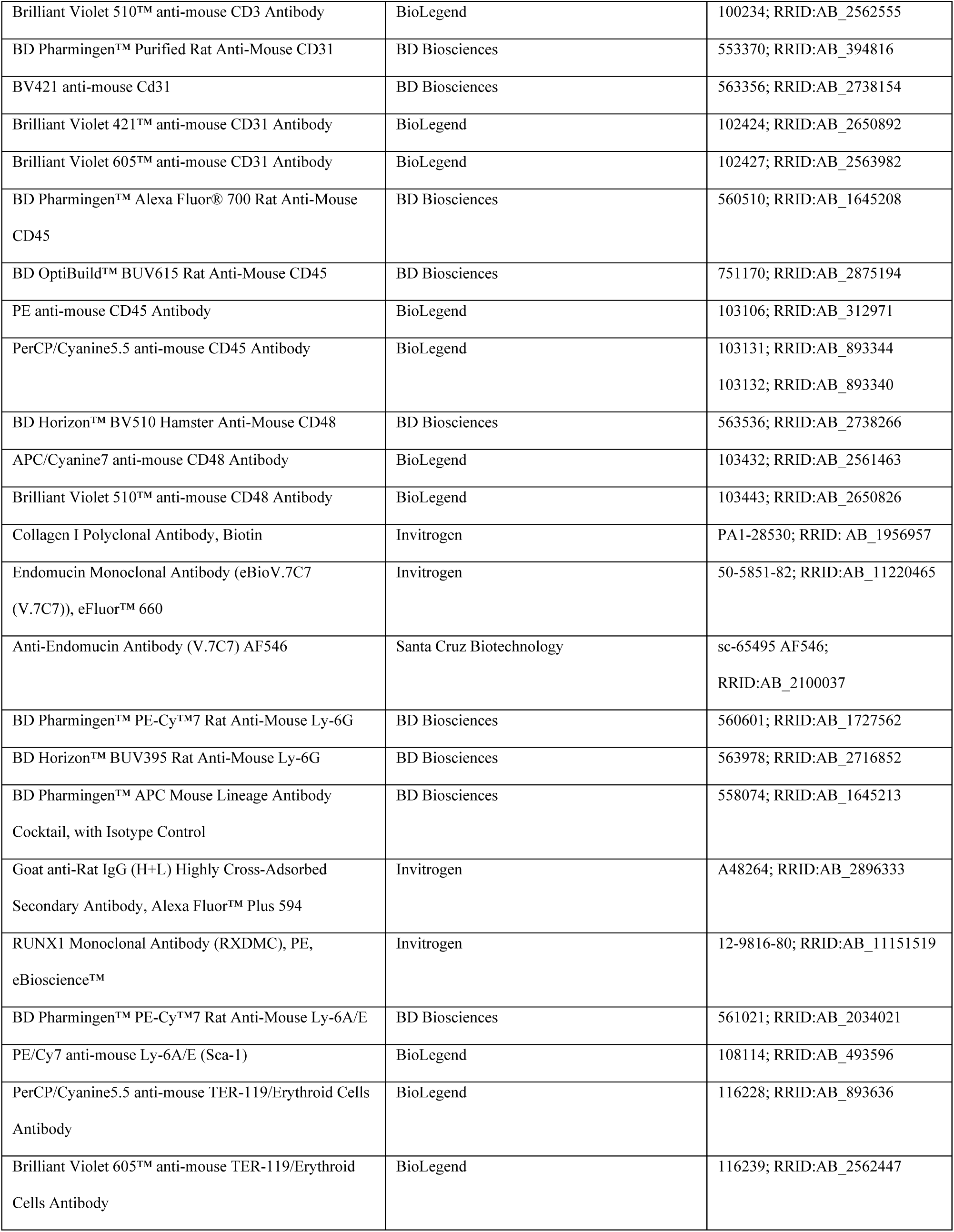
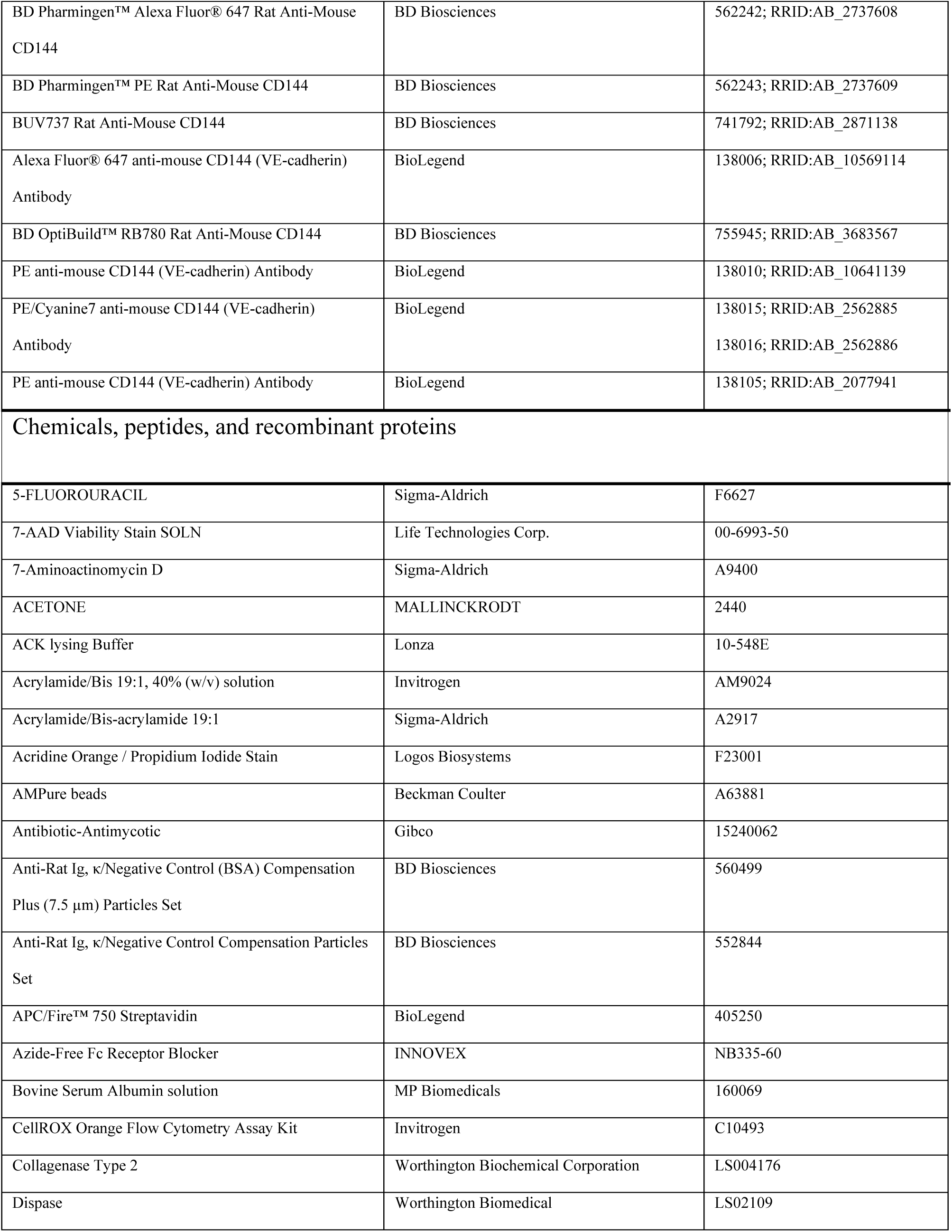
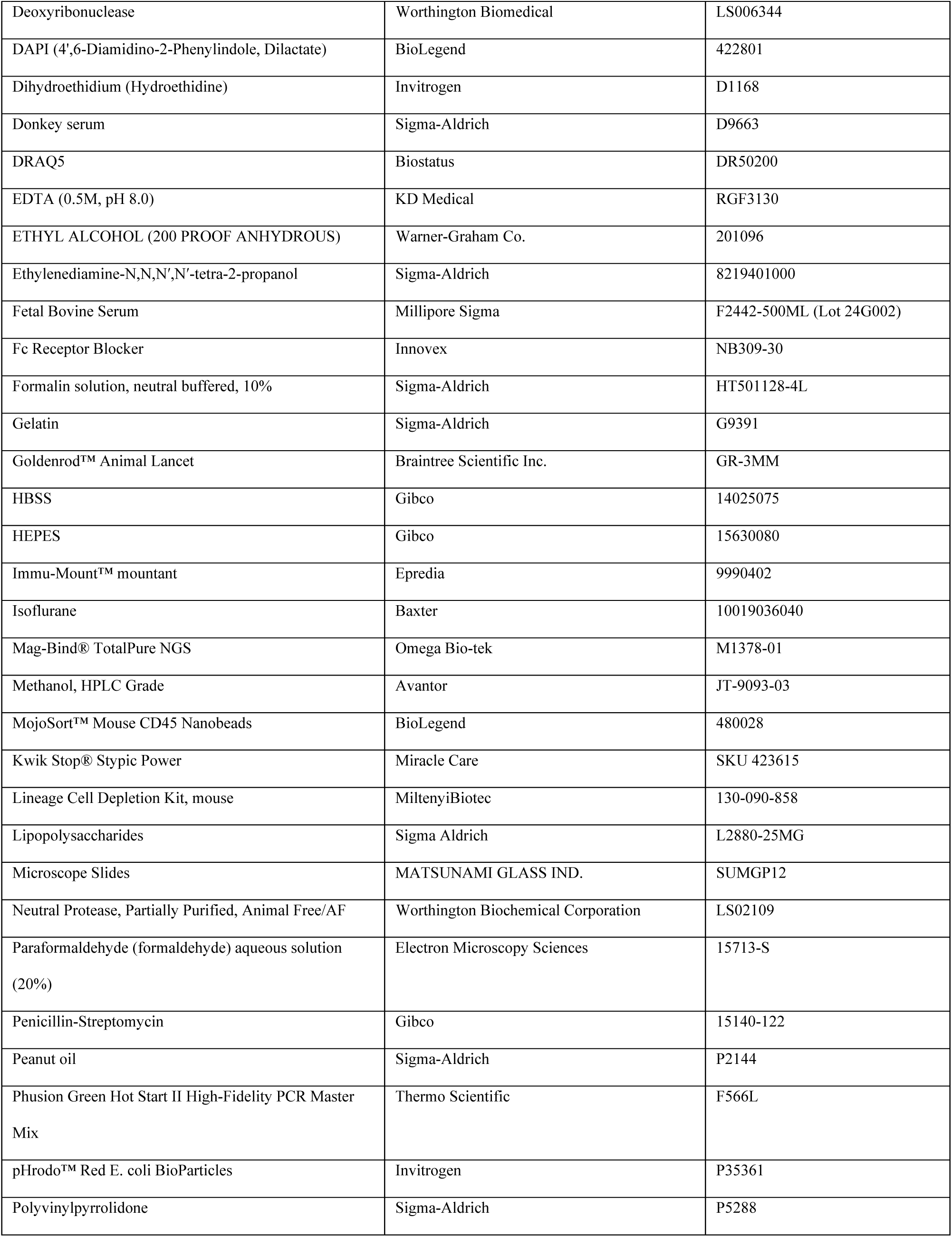
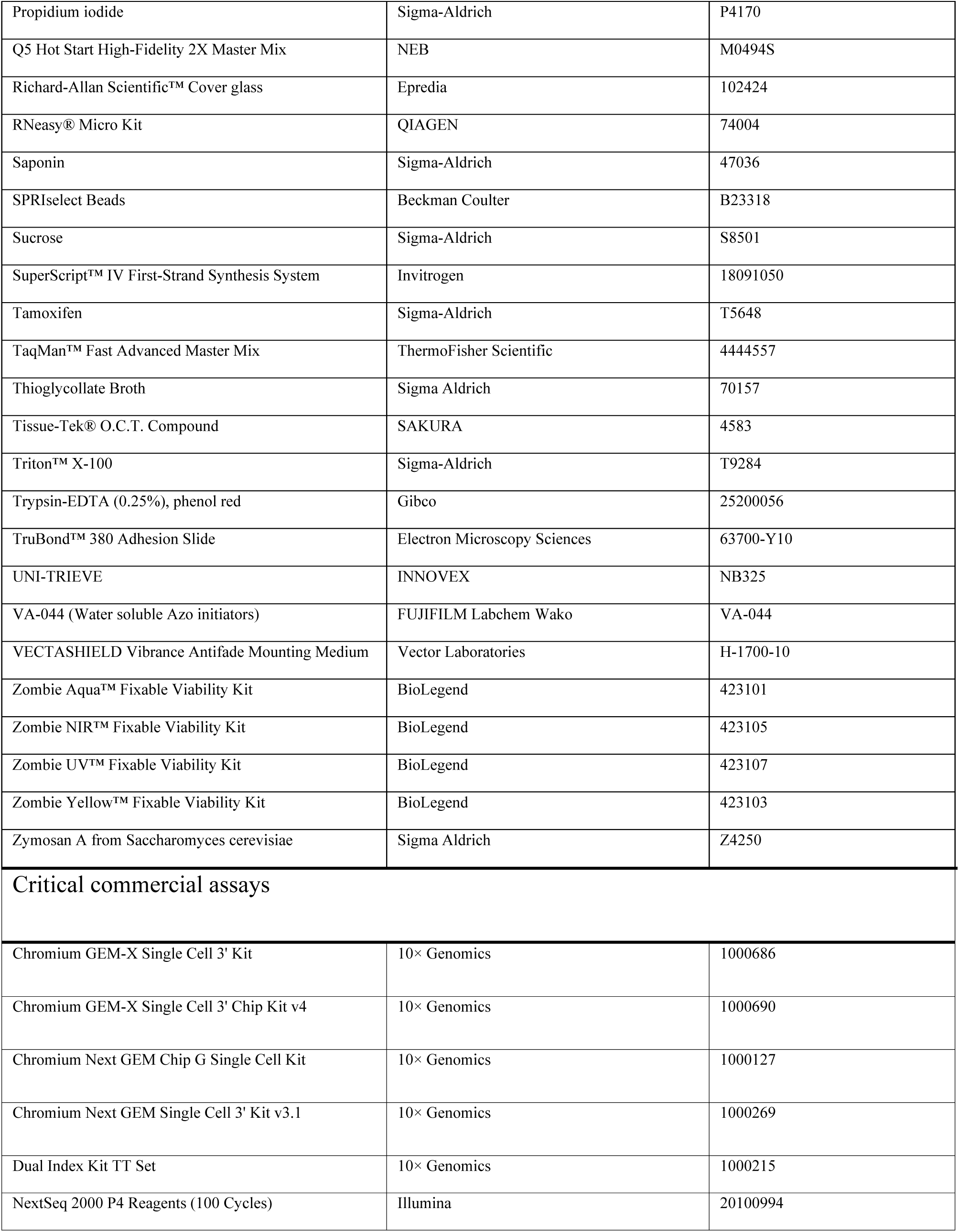
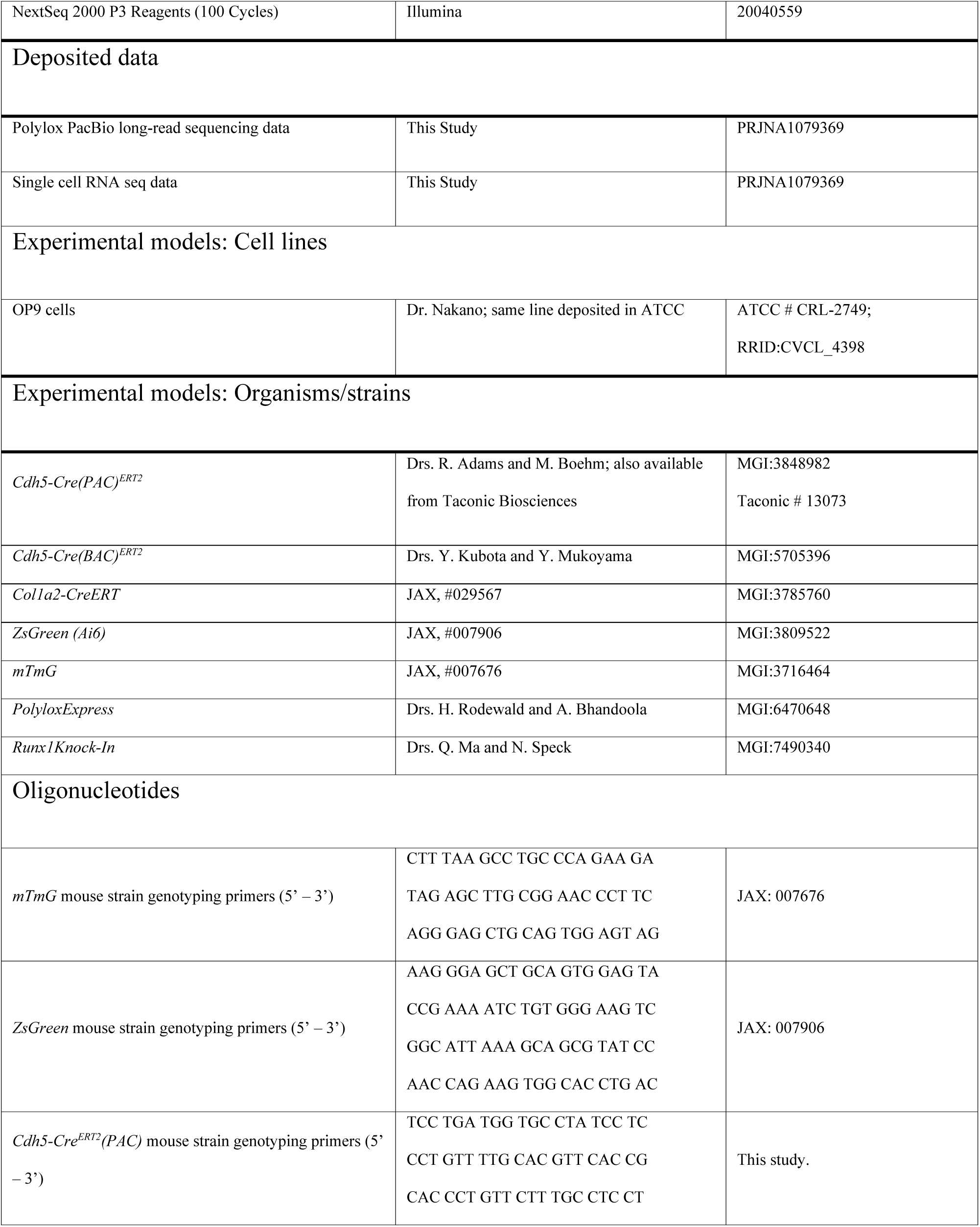
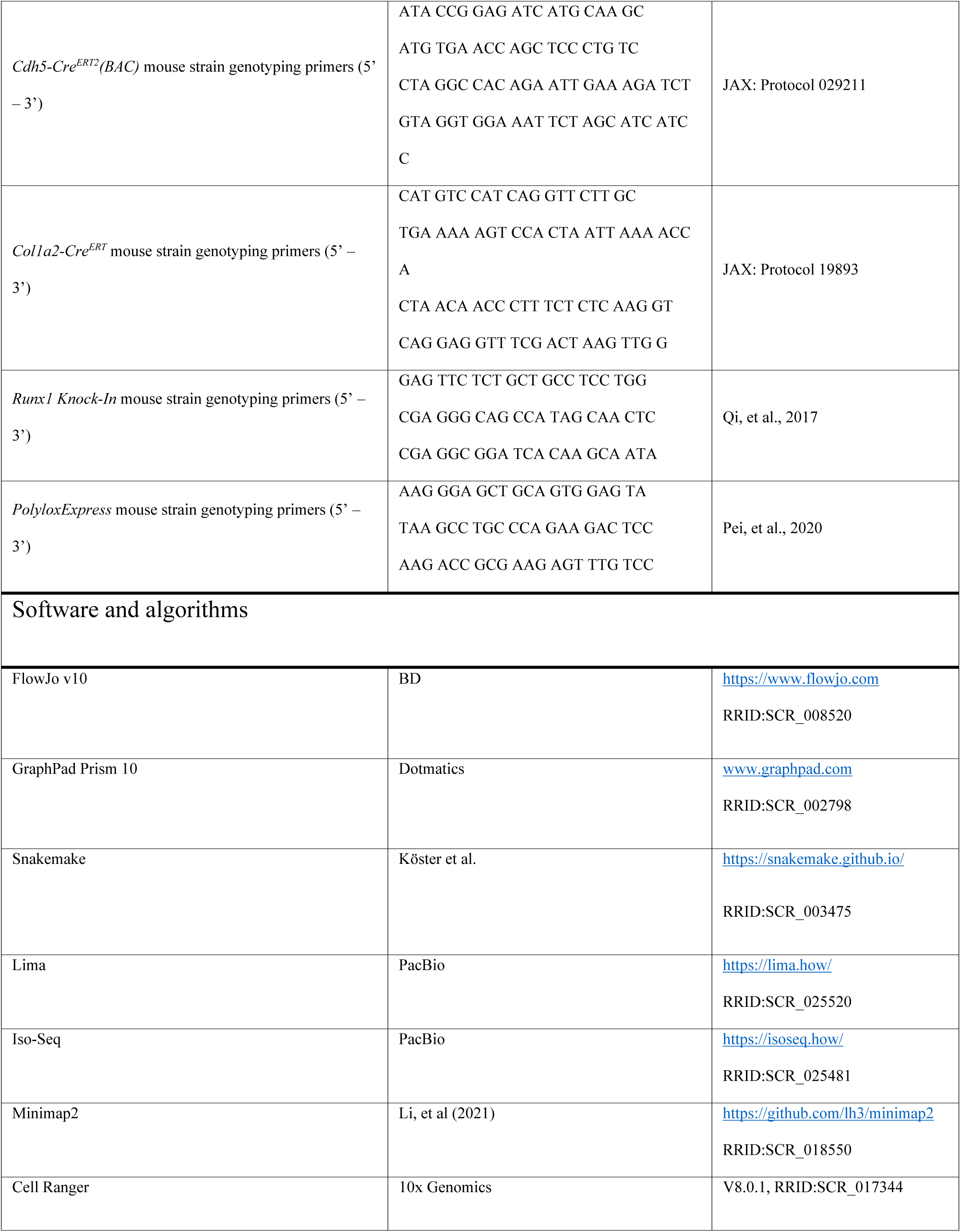
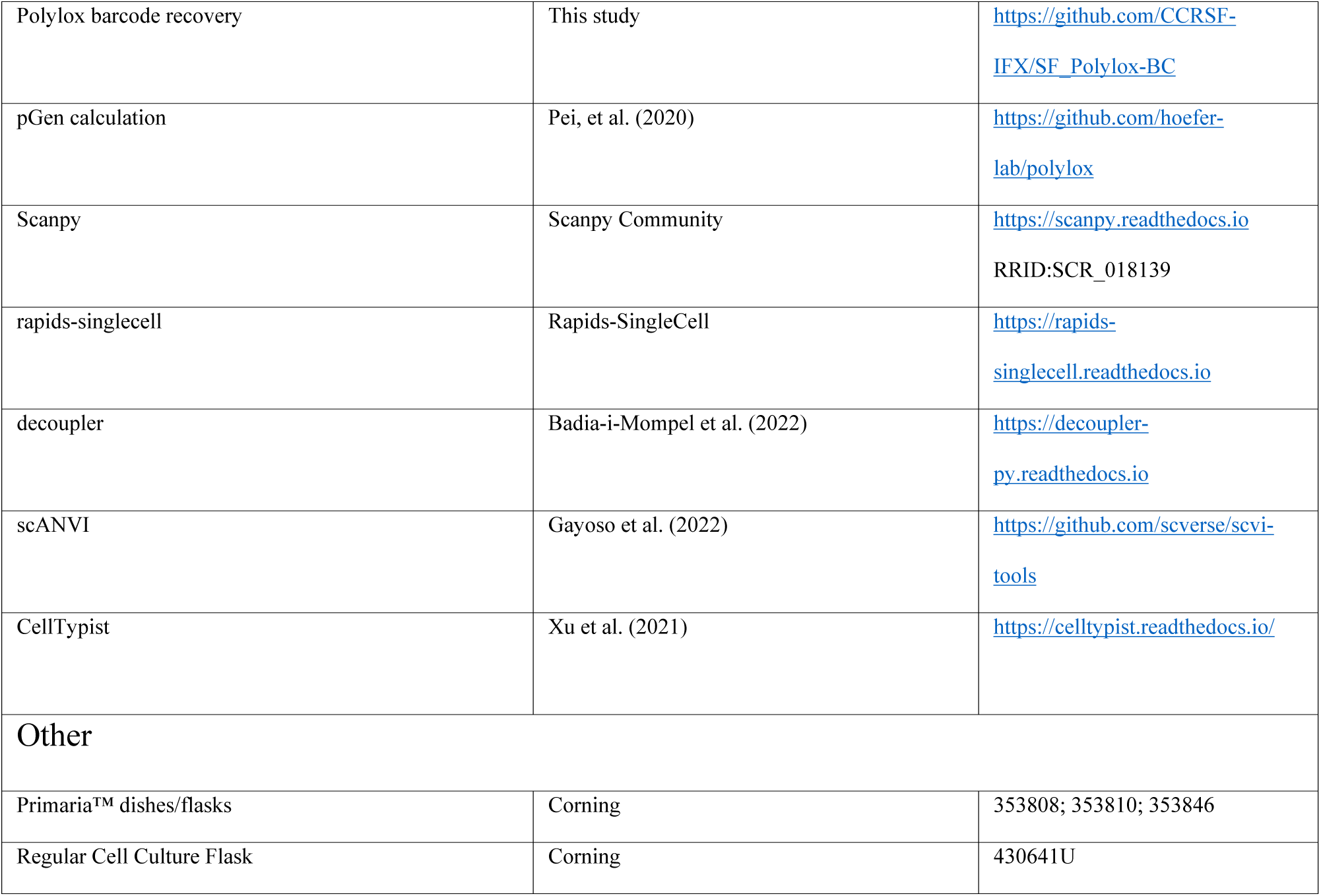

### EXPERIMENTAL MODEL AND STUDY PARTICIPANT DETAILS

#### Mouse strains

All animal studies were approved by the Institutional Animal Care and Use Committee of the CCR (Bethesda, MD), National Cancer Institute (NCI), NIH and conducted in adherence to the NIH Guide for the Care and Use of Laboratory Animals (National Academies Press, 2011) and approved protocols. *Cdh5-Cre^ERT2^(PAC)* mice ^37^ (MGI:3848982) were a gift from Dr. R. Adams and *Cdh5-Cre^ERT2^(BAC)* mice ^39^ (MGI:5705396) were a gift from Dr. Kubota. *Col1a2-CreER* mice ^85^ were purchased from the Jackson Laboratory (JAX#029567). Cre-dependent *Ai6-ZsGreen* ^86^ (JAX#007906) and *mTmG* ^87^ (JAX#007676) fluorescent reporter mice were purchased from the Jackson Laboratory. In *ZsGreen* reporter mice, Cre activity leads to constitutive expression of ZsGreen1 in cell bodies. In *mTmG* reporter mice, Cre activity leads to an irreversible switch from cell membrane-localized tdTomato (mT) to membrane-localized EGFP (mG). *PolyloxExpress* mice ^33^ were a kind gift of Drs. Hans-Reimer Rodewald and Avinash Bhandoola. The Cdh5-Cre^ERT2^/ZsGreen/*PolyloxExpress* mouse line was generated in house. Runx1 conditional knock-in mice^66^ (Gt(ROSA)26Sor^tm1(CAG-Runx1)Lzjg^, MGI:7490340) were generously provided by Drs. Qiufu Ma and Nancy Speck. Runx1 endothelial-specific knock-in (*Runx1^EC-KI^*) mice were generated by crossing *Runx1^Ki/+^* mice with *Cdh5-Cre^ERT2^(PAC)/ZsGreen^Tg/Tg^* mice. Upon tamoxifen treatment, Cre-mediated excision of the two floxed STOP codons enables co-expression of Runx1 and ZsGreen in ECs.

All animals were bred in the animal facilities of CCR/NCI (Bethesda, MD). The mice were maintained in a C57BL/6J background. Mice were identified with ear tags and routinely genotyped by PCR. No mouse was excluded from the experiments, unless assessed as sick by the veterinarians or fight wounds were observed at harvest. Tamoxifen (Sigma-Aldrich, #T5648) dissolved in peanut oil (Sigma-Aldrich, #P2144) (10 mg·mL^-1^) was administered orally (via gavage using 22g feeding needles) at 100 mg·kg^-1^. Unless otherwise specified, three doses on consecutive days were administered. Unless indicated otherwise, 8- to 12-week-old male and female mice were used when tamoxifen was administrated. No randomization or blinding was used to allocate experimental groups. Mice were typically sacrificed between 9 AM and 11 AM local time.

## METHOD DETAILS

### OP9 cell culture

OP9 cells, a gift from Dr. T. Nakano^44^ (also available from ATCC, #CRL-2749), were maintained in α-MEM (Gibco #12561056; without ribonucleosides and deoxyribonucleosides, with 2.2 g/L sodium bicarbonate) supplemented with 20% fetal bovine serum (FBS; Sigma-Aldrich #2442). Culture dishes and flasks (Corning #353003, #430641U) were precoated with gelatin (Sigma-Aldrich #G9391; ∼100 µL/cm², 60 minutes at 37°C). Cells were incubated at 37°C in a humidified atmosphere of 95% air and 5% CO₂.

### Bone marrow cell harvesting, culture and terminal harvest for analysis

To harvest BM cells to be cultured, mice were euthanized by cervical dislocation and soaked in 70% ethanol for 5 minutes. Long bones (femurs and tibiae) were dissected, and surrounding skin, muscle, and connective tissue were carefully removed. Cleaned bones were immediately transferred into ice-cold sterile PBS and kept on ice. Once all bones were harvested, they were soaked in 70% ethanol for 1 minute and rinsed three times with ice-cold PBS.

Each bone was cut into two pieces using sterile scissors and placed--with the open end facing downward--into a sterile 500 µL microcentrifuge tube pre-perforated with a 16G needle (sterilized in advance). Each 500 µL tube was loaded with two femurs and two tibiae. To each tube, 200 µL of DMEM supplemented with 1 mM EDTA was added. The 500 µL tubes were then placed inside sterile 1.5 mL microcentrifuge tubes, sealed with Parafilm, and centrifuged at 12,000 rpm for 20 seconds. The inner tubes were discarded, and the BM pellet collected in the 1.5 mL tubes was resuspended in 1 mL DMEM + 1 mM EDTA.

Cells were filtered through 70 µm cell strainers, centrifuged at 350 × g for 5 minutes at 4°C, and resuspended in complete DMEM (DMEM containing 15% FBS (Sigma-Aldrich #2442-500ML, LOT:24G002), Penicillin-Streptomycin [Gibco #15140-122], and Anti-Anti [Gibco #15240-062]). A single-cell suspension was prepared by pipetting 30 times with a 1 mL pipette.

Bone marrow cells were cultured either on Corning Primaria™ dishes/flasks (Corning #353808, #353810, #353846) or on OP9 stromal cell monolayers in standard tissue culture dishes/flasks (Corning #353003, #430641U) precoated with gelatin (Sigma-Aldrich #G9391; ∼100 µL/cm², incubated 60 min at 37°C).

Cells were seeded at a density equivalent to BM cells from two femurs and two tibiae per ∼75 cm² culture surface (e.g., T75 flask) in 15 mL complete DMEM. After ∼32 hours, non-adherent cells and medium were removed, and adherent cells were gently washed three times with PBS. For Primaria cultures, 15 mL fresh complete DMEM was replenished twice weekly. For OP9 co-cultures, non-adherent cells and medium were removed twice weekly and replaced with fresh complete DMEM supplemented with freshly isolated unfractionated WT BM cells (from two femurs and two tibiae per ∼75 cm² surface, using the same BM isolation method mentioned above).

For terminal collection, at the start of the final culture week, all non-adherent cells were removed, and adherent cells were washed three times with PBS. Fresh complete DMEM (15 mL) was added. After three days, a second 15 mL complete DMEM addition was made. No further medium changes occurred before harvest. For final collection, non-adherent cells and supernatant were collected first. Adherent cells were washed with PBS, and the wash was pooled with the supernatant. Remaining adherent cells (including OP9 and BM-derived cells) were detached using 0.25% Trypsin-EDTA (Gibco #25200-056) for 5 minutes at 37°C and added to the pooled suspension. After another PBS wash, a second trypsinization (10 minutes at 37°C) was performed. All collected material from each step was combined for downstream flow cytometric analysis. For transplantation experiments, only non-adherent cells (with supernatant) and loosely adherent cells recovered after the initial 5-minute Trypsin-EDTA incubation were pooled and subjected to FACS sorting to isolate ZsGreen⁺ cells for injection.

### Blood collection

For terminal collection, blood was obtained from the mouse abdominal aorta with BD Vacutainer™ EDTA tubes (BD #367856) and Vaculet™ blood collection needles (23G, EXELINT #26766). Blood smears were prepared with 10 µL of collected blood. For flow cytometry analysis, ACK buffer (Lonza, #BP10-548E) was added to the blood to lyse red blood cells before Fc receptor blocking and antibody staining. For non-terminal blood collection, ∼20-50µL blood was collected by submandibular blood sampling, using a 3mm animal lancet (BRAINTREE SCI., GR-3MM) and a 250µL BD Microtainer^®^ K2EDTA tube (BD #365974).

Kwik Stop^®^ Styptic Powder was applied to stop the bleeding (Miracle Corp. #423615). For long-term, repeated blood collection, 2 drops of blood were collected from the tail vein with Microhematocrit Capillary Tubes (Fisherbrand # 22-362574). White blood cell (WBC) counts were determined using acridine orange/propidium iodide (AO/PI, Logos Biosystems, #F23001) staining and quantified with a LUNA-FL™ fluorescence cell counter (Logos Biosystems). For flow cytometry detection of ZsGreen/EGFP positive PBMCs, blood was collected with one of the methods above. For flow cytometric detection of ZsGreen⁺/EGFP⁺ PBMCs, blood was collected using one of the three methods described above. Red blood cells were lysed with ACK buffer, and DAPI and DRAQ5 were used to exclude dead cells and to identify nucleated PBMCs.

### Bone marrow harvest for flow cytometry analysis

For flow cytometry analysis, bone marrow was harvested using one of the two methods described below. For hematopoietic cell isolation, bone marrow was harvested by flushing femurs and tibiae with ice-cold Sort Buffer (1× PBS [Gibco, #10010-031] supplemented with 5 mM EDTA, 25 mM HEPES, and 2% FBS [Sigma-Aldrich, #F2442]). Red blood cells were then lysed using ACK lysing buffer (Lonza, #10-548E) according to the manufacturer’s instructions. Cells were then washed with Sort Buffer and passed through a 40μm cell strainer (GREINER BIO-ONE, #542040, #542140). For greater preservation of endothelial cells, bone marrows were harvested by gently crushing mouse femurs and tibiae in Sort Buffer (5mM EDTA, 25mM HEPES, 2% FBS in 1× PBS). Red cell lysis was performed using ACK buffer. Bone marrow cells were then incubated with 0.1U·mL^-1^ Collagenase (Worthington Biomedical Corp., #LS004176), 0.8U·mL^-1^ Dispase (Worthington Biomedical Corp., #LS02109), and 0.5mg·mL^-1^ DNase (Worthington Biomedical Corp., #LS006344) in 1x Hanks’ Balanced Salt Solution (HBSS) with Ca^2+^ and Mg^2+^ (Gibco, # 14065056) at 37°C for 30 min on a rotating mixer. Cells were then washed with Sort Buffer and passed through a 40μm cell strainer.

### Endothelial cell transplantation experiments

Four days prior to transplant, recipient mice received one dose of 5-FU (150 mg·kg^-1^ in PBS, Sigma-Aldrich, #F6627) intraperitoneally under isoflurane anesthesia. Sorted bone marrow cells (Col1a2-Cre/ZsGreen: 5,000 cells; Cdh5-Cre/ZsGreen: 20,000 cells in 100μL PBS) were inoculated retro-orbitally under isoflurane anesthesia. Bone marrows and blood were harvested from transplant recipients four weeks after the transplant unless otherwise specified.

### Bone marrow ablation and transplantation

Recipient mice were lethally irradiated (11 Gy) using a Cesium-137 (¹³⁷Cs) gamma irradiator three days prior to transplantation.

For whole BM cell transplantation into *mTmG* recipients, BM cells were harvested from WT donor mice by flushing (as described above); 5 × 10^6^ cells were transplanted per recipient via tail vein. No rescue bone marrow cells were infused.

For LSK transplantation into WT recipients, BM from Cdh5-Cre^ERT2^(PAC)/ZsGreen mice (not treated with tamoxifen) was harvested by flushing, followed by red blood cell lysis with ACK buffer. Fc receptor blocking was performed using Azide-Free Fc Receptor Blocker (Innovex, #NB335-60) per manufacturer’s instructions. Cells were stained with antibodies to Lineage cocktail, Sca-1, and c-Kit. DAPI was used to exclude dead cells. LSK populations were sorted using Sony SH800S and its Ultra Purity mode. Transplanted cell numbers were as follows: ZsGreen⁻ LSKs (5 × 10^4^, n = 3; 2.5 × 10^4^, n = 3) and ZsGreen⁺-enriched LSKs (2,800 cells, n = 2). No rescue bone marrow cells were infused.

For transplantation of ex vivo–cultured BM cells, non-adherent and loosely adherent ZsGreen⁺ cells were collected from 8-week OP9 co-cultures (as described above), sorted by FACS for live (DAPI^-^) ZsGreen^+^ cells, and transplanted into irradiated recipients at 5 × 10^4^, 2.5 × 10^4^, or 1.25 × 10^4^ cells per mouse (n = 2 per group). No rescue bone marrow cells were infused.

### Flow cytometry and cell sorting

For intracellular antigen detection, single cell suspensions of bone marrow and blood were first incubated with Azide-Free Fc Receptor Blocker (Innovex, #NB335-60), following the manufacturer’s instructions. After washing, cells were first stained with surface marker antibodies at the concentration of 2μg per 1×10^7^ cells in Sort Buffer for 30 minutes at 4°C and then stained with live/dead cell discriminating BioLegend Zombie Dyes (UV, NIR, Violet, or Yellow, BioLegend #423108, #423106, #423114, and # 423104) following the manufacturer’s instructions. After washing, cells were fixed in 4% paraformaldehyde for 10 minutes at 37°C, and then permeabilized with 1% saponin (Sigma-Aldrich, # 47036) /Sort Buffer for 30min on ice. Subsequently, the cells were stained with rat monoclonal primary RUNX1-PE Ab (Invitrogen, # 12-9816-80) in 0.1% saponin/Sort Buffer overnight. After washing and resuspension, propidium iodide (PI, 0.5μM, Millipore Sigma, # P4170), 7-AAD (1μg·mL^-1^, Millipore Sigma, #A9400) or DAPI (0.5μg·mL^-1^, BioLegend, #422801) was added as a nuclear counterstain. For live cell staining without cell permeabilization, after cell surface antibody staining, cells were washed and suspended in Sort buffer containing propidium iodide (PI, 0.5μM, Millipore Sigma, # P4170), 7-AAD (1μg·mL^-1^, Millipore Sigma, #A9400) or DAPI (0.5μg·mL^-1^, BioLegend, # 422801), to distinguish dead cells from the live cells. For FACS sorting of live endothelial cells, after Fc receptor blocking, bone marrow cells were first depleted of CD45^+^ cells with MojoSort mouse CD45 nanobeads (BioLegend #480028) following the manufacturer’s protocol and then stained with specific antibodies. For flow cytometry analysis, compensation beads (BD Biosciences, #552844) were used for flow cytometer compensation.

Flow cytometric data were acquired with BD FACSCanto-II, BD LSRFortessa, BD FACSymphony A5 (BD Biosciences), Sony SA3800 or Sony ID7000 cell analyzers. Cell sorting was performed with BD FACS Aria III, BD FACS Aria Fusion or Sony SH800S cell sorters.

FSC and SSC profiles were used for excluding dead cells and debris. 7-AAD, PI, DAPI or BioLegend Zombie Dye was used for excluding dead cells. FSC-W versus FSC-H and SSC-W versus SSC-H were used to gate on single cells. Unless otherwise specified, fluorescence minus one (FMO) controls are used for negative gating reference. For BM HSPC analysis, BM cells were harvested followed by lineage positive cell depletion (Biolegend #480004). Data were analyzed with FlowJo (BD, v10.8.1), SONY ID7000 Software (Version 1.2.0.28212) or FACS Diva (BD, v6.1 and v9.0). Flowjo Plugins UMAP_R (v4.0.4) and FlowSOM (v4.1.0) were used for UMAP dimensional reduction and unsupervised clustering of flow cytometry data. Following sorting, a small aliquot of collected events were re-run on the sorter to assess purity. Analysis was performed using the identical gating hierarchy (FSC/SSC → singlets → live cells → target population).

### Bone marrow cryosections

Deeply anesthetized mice were transcardially perfused with 20mL ice-cold 1x PBS, followed by perfusion with 15mL ice-cold hydrogel solution (5% acrylamide/bis-acrylamide 19:1 (Sigma-Aldrich, #A2917), 2.5mg/mL polymerization initiator VA-044 (FUJIFILM, Wako, VA-044, Water soluble Azo initiators), 4% PFA in 1× PBS, 5mL/min flow speed). Femurs and tibiae were collected in tubes containing 5mL hydrogel solution and incubated at 4°C for 4 hours. The bones were then washed with PBS and incubated at 37°C for 2 hours. Bone decalcification was performed by incubating the bones in 40mL 0.5M EDTA pH 8.0 (KD Medical, #RGF-3130) for 3 days on a rotate mixer, with daily refreshed 0.5M EDTA solution. The bones were then dehydrated in 20% sucrose and 2% polyvinylpyrrolidone in PBS overnight. Dehydrated bones were then embedded in OCT (SAKURA, #4583) blocks using Precision Cryoembedding System (IHC WORLD, #IW-P101). Cryosections (10μm) for immunofluorescence staining were prepared from OCT frozen bone blocks using Leica CM3050S microtome, low-profile microtome blades (Leica 819, #14035838925), and TruBond™ 380 adhesion slides (Electron Microscopy Sciences, # 63701-W10).

### Immunofluorescence staining and imaging

Tissue sections were rehydrated with 1× PBS (15 minutes), permeabilized in 0.3% Triton X-100 (Sigma-Aldrich, #T9284)/PBS (15 minutes), washed in 1× PBS, and incubated (2 hours) with blocking solution (2% BSA, 5% donkey serum (SIGMA, #D9663), and 0.3% Triton X-100/PBS). Samples were then washed 3 times with PBS and incubated with primary antibodies (5ng/mL; 4°C overnight). When secondary antibodies were used, 3 PBS washes were performed before incubating with fluorescent-labeled secondary antibodies (2ng/mL, room temperature, 2 hours). After washing (3x, 10 minutes each with 1× PBS), DAPI was added (300nM in PBS, 10 min). After 3 washes (5 min each with 1× PBS), coverslips were mounted (EPREDIA, #9990402), dried and sealed with nail polish. For blood smear staining, slides were first soaked in acetone/methanol/PFA (19:19:2 for 90 seconds) before rehydration ^88^. Confocal imaging was performed with Zeiss LSM 780, Zeiss LSM 880 NLO Two Photon, or Nikon *ECLIPSE* Ti2-E SoRa systems, according to the experimental specific needs (resolution, speed, wavelength capabilities). Images were processed with Zen (Zen Black v2.3, release Version 14.0.12.201, Zen Blue Lite v2.5, Carl Zeiss), Bitplane Imaris (v9.7.0, Oxford instruments), and Photoshop (v23.3.0, Adobe, for whole image contrast and brightness adjustments).

### Isolation of peritoneal cavity cells

To isolate peritoneal cavity cells, mice were euthanized by cervical dislocation, injected intraperitoneally (i.p.) with cold FACS sort buffer (5 mL), massaged and flicked gently on the abdomen. Peritoneal fluid was then withdrawn slowly and transferred into a polypropylene centrifuge tube on ice prior to centrifugation (350×g, 10min, at 4°C) and analysis.

### Thioglycolate induced sterile peritonitis

Mice were injected i.p. with 1 mL PBS or 4% thioglycolate (Sigma-Aldrich, #70157). Peritoneal cavity cells were harvested 4 hours after injection for further analysis.

### Phagocytosis assay

Phagocytosis assay was performed using pHrodo™ Red *Escherichia coli* BioParticles (Invitrogen, # P35361) following the manufacturer’s instructions. Briefly, mice were first treated with thioglycolate (as described). Peritoneal cavity cells were incubated with the fluorescent *Escherichia coli* particles for 60 min at 37°C. Cells were then stained with surface marker antibodies and analyzed by flow cytometry.

### ROS assay

ROS assay was performed using CellROX™ Orange Flow Cytometry Assay Kit (Invitrogen, # C10493) following the manufacturer’s instructions. Briefly, mice were first treated with thioglycolate (as described). CellROX detection reagent was added to peritoneal cavity cells (final concentration of 500nM) prior to incubation for 60 min at 37°C. Cells were then stained with the surface marker antibodies and analyzed by flow cytometry.

### PolyloxExpress single cell lineage tracing

These experiments essentially followed published procedures ^33^. Briefly, BM cells from tamoxifen treated Cdh5-Cre/ZsGreen/Polylox mice (treated at 10 weeks old and harvested at 16 weeks old) were enriched for ZsGreen positive cells by FACS. Single cell capturing was performed with 10x Genomics Chromium Single Cell 3′ Reagent Kits, following manufacturer’s protocols. After reverse transcription (RT) in droplets, pooled cDNA was amplified and split into two aliquots for parallel transcriptome library preparation and barcode enrichment. Initial library quality control was performed with Agilent TapeStation D5000. For transcriptome analysis, 10 μL (25%) of a 10x cDNA library was fragmented and a gene expression library, generated following protocols in Single Cell 3’ Reagent Kits v3 and v4, was sequenced with Illumina NextSeq 2000 P3/P4 Reagents (28+74 bp read length). For Polylox barcode amplification, targeted amplification of barcodes from a 5-10 ng aliquot of a 10x cDNA library was performed by nested PCR. In the first round, primers #2,652 (5′-GCATGGACGAGCTGTACAAG-3′, annealing at the 5′ end of Polylox) and #2,674 (5′-AATGATACGGCGACCACCGAGATCTACACTCTTTCCCTACACGACGCTC-3′, annealing at the adaptor site (read 1) were used for amplification for 5 min at 95°C; (30 s at 95°C, 30 s at 57°C, 3 min at 72°C) 12 times; 10 min at 72°C. PCR products purified with 0.7x AMPure beads were used for the second round of PCR using primers #2,426 (5′-CGACGACACTGCCAAAGATTTC-3′, annealing at the 5′ end of Polylox) and #2,676 (5′-AATGATACGGCGACCACCGA-3′, annealing at the 5′ end of primer #2,674), for 5 min at 95°C; (30 s at 95°C, 30 s at 60°C, 3 min at 72°C) 18 times; 10 min at 72°C. The PCR products were purified with AMPure PB beads according to the manufacturer’s protocol. Long read amplicon-seq libraries were sequenced by PacBio Sequel II with PacBio Amplicons Library Preparation using SMRTbell prep kit 3.0. A custom transcriptome reference was built from mouse reference MM10 to include ZsGreen1.

A snakemake workflow with custom python script was used to retrieve cell indexes and Polylox barcodes from the PolyloxExpress amplicons (https://github.com/CCRSF-IFX/SF_Polylox-BC).

Single cell transcriptome and single cell barcodes were linked using the 10x 3’ kit cell index and group information. Further analysis and illustrations were generated using Scanpy (https://github.com/scverse/scanpy). Doublet detection was performed using Scrublet, with the predicted doublet rate calculated based on 10x Genomics guidance about the cell numbers loaded to the microfluidic chips. Batch correction was done using BBKNN method (https://github.com/Teichlab/bbknn). Cell types were manually annotated based on canonical marker gene expression, guided by results from three automated annotation tools: (1) decoupleR ((https://saezlab.github.io/decoupleR/), using PangLaoDB (https://panglaodb.se/) as reference; (2) scANVI (https://github.com/scverse/scvi-tools) using ImmGne (https://www.immgen.org/) as reference; (3) CellTypist, using the embedded Immune_All_Low model. The rare barcodes and their pGen were identified using the MatLab script from the Höefer’s Lab (https://github.com/hoefer-lab/polylox). True barcodes are defined as pGen < 1×10^-4^, such that the expectation of a True barcode in detected 4,072 cells is 0.407. This work utilized the computational resources of the NIH HPC Biowulf cluster (http://hpc.nih.gov) and Frederick Research Computing Environment (FRCE). Python version: 3.10.; R version 4.5.0.

### Public single-cell RNA-seq data analysis

Raw FASTQ and BAM files were downloaded from publicly available datasets (GSE108885 ^54^, GSE108891 ^54^, GSE118436 ^54^, GSE123078 ^54^, GSE122465 ^55^, GSE128423 ^53^, GSE145477 ^65^, GSE156635 ^56^, GSE137116 ^24^, E-MTAB-8077 ^65^, GSE230260^58^, GSE259382^59^). The BAM files were first converted to FASTQ files with bamtofastq (10x Genomics, v2.0.1). All FASTQ files were then processed by the count function of Cell Ranger (10x Genomics, v8.0.1) and aligned to the mouse genome (mm10, version 2020-A), to generate read matrixes. Further analysis and illustrations were generated using Scanpy as described above.

### Definition of cultured fluorescent BM cell clusters and quantification

A cluster was defined as (1) a spatially distinct group of ZsGreen⁺ cells not contiguous with other fluorescent cells, (2) containing a central core with at least five extending branches, and (3) exhibiting a roughly radial organization, with projections spreading outward in multiple directions, and (4) extending at least ∼200 µm in one direction. Isolated or irregularly scattered fluorescent cells were not counted. Due to their large size, clusters were counted manually across the culture dish using a tally counter under fluorescence microscopy.

### RNA isolation and quantitative RT PCR

RNA was extracted using RNeasy^®^ Micro Kit (QIAGEN, #74004), following the manufacturer’s protocol. Cells were sorted directly into RNA lysis buffer (Buffer RTL of RNeasy^®^ Micro Kit). cDNA samples were prepared with SuperScript IV Reverse Transcriptase (Invitrogen, #18091050), following manufacturer’s instructions. Real Time PCR was performed using Applied Biosystems™ TaqMan™ Fast Advanced Master Mix (4444557) and Applied Biosystem™ QuantStudio™ 5 Real-Time PCR System. TaqMan™ probes used were purchased from Applied Biosystems™: Spp1 (Mm00436767_m1); Cxcl12 (Mm00445553_m1); Col1a2 (Mm00483888_m1); Cdh5 (Mm00486938_m1); Runx1 (Mm01213404_m1); Ptprc (Mm01293577_m1); Gapdh (Mm99999915_g1); Actb (Mm02619580_g1). The reaction condition was set as follows: 50°C 2 minutes, 95°C 20 seconds, 45 cycles of 95°C 1 second, 60°C 20 seconds. Ct values were determined using the ABI QuantStudio™ Design & Analysis Software (v1.5.2). Relative gene expression was assessed using the 2^-ΔΔCt^ method, normalized to Gapdh expression level for each sample. The data was further normalized to gene expression levels in the unsorted bone marrow sample to calculate relative gene expression levels in each sample. Data reflect triplicates real-time PRC experiments.

## QUANTIFICATION AND STATISTICAL ANALYSIS

No statistical method was used to predetermine sample size. No data were excluded from the analyses. Mice with the correct genotypes were randomly assigned to control or treated groups. Unless otherwise specified, data are represented as mean ± S.D, and individual dots in the graphs indicate individual mice. Comparisons between two groups were performed using two-tailed unpaired Student’s *t*-tests (except for Figure S1-H, which used a paired *t*-test). Spearman rank correlation test was used for Figure 3E. Statistical analyses were performed with GraphPad Prism (v9.0.1). A statistical difference of *P*<0.05 was considered significant: ns, not significant, * *P* < 0.05, ** *P* < 0.01, *** *P*< 0.001.

## Supplemental Figure Legends

**Figure S1.**
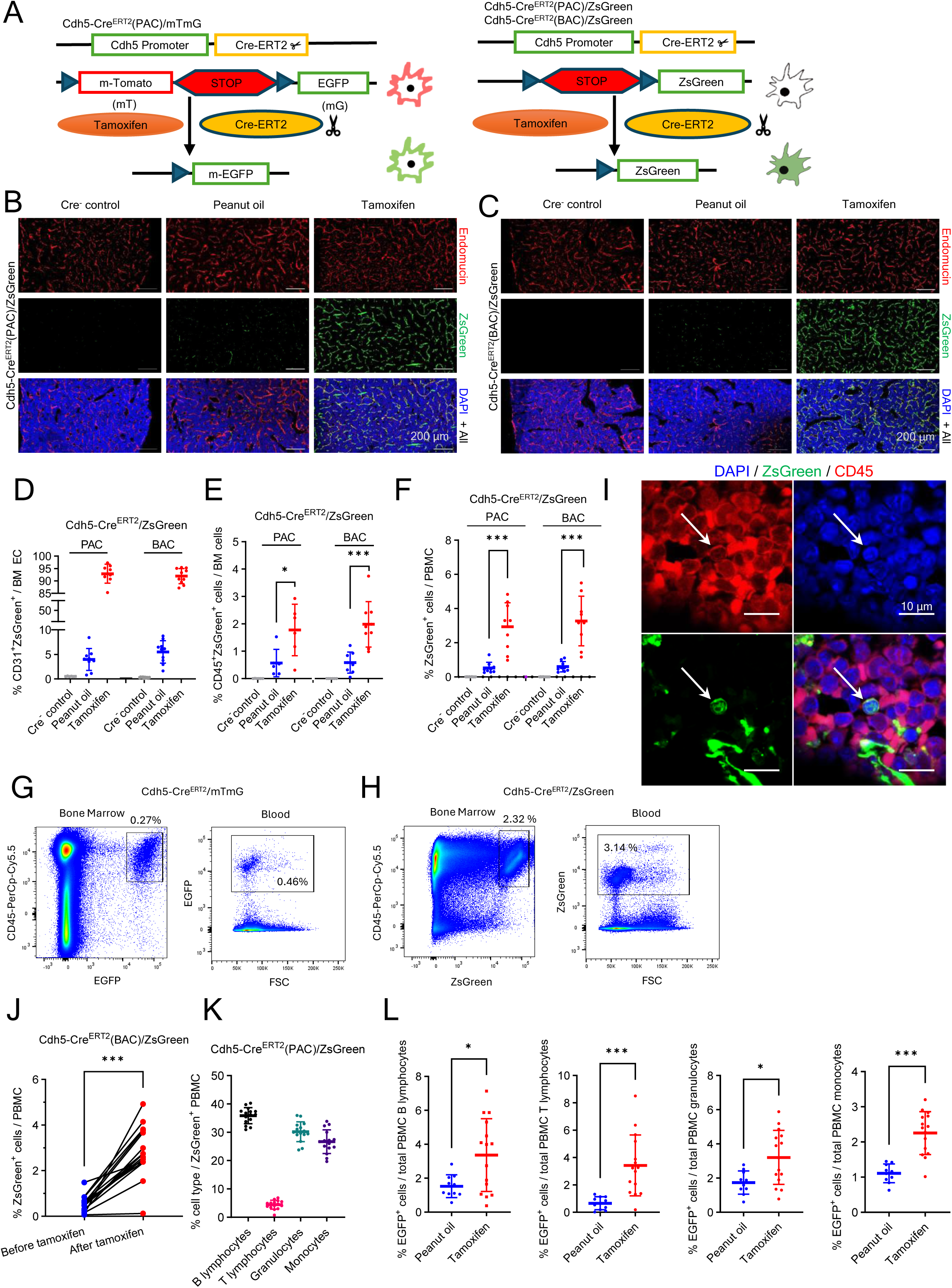
Contribution of ECs to hematopoiesis in adult BM is revealed by Cdh5-Cre^ERT2^ mouse tracking lines. Related to Figure 1. (A) Cdh5-tracking mouse lines. Tamoxifen switches-on green fluorescence in cells that express the Cre-recombinase and their cell progeny. (B and C) Confocal microscopy images of representative BM sections from Cdh5-Cre^ERT2^(PAC)/ZsGreen (B) and Cdh5-Cre^ERT2^(BAC)/ZsGreen (C) adult mice showing tamoxifen-induced ZsGreen fluorescence co-staining of most Endomucin^+^ cells. Control BM sections from representative Cre^+^ mouse treated with peanut oil (no tamoxifen) display occasional ZsGreen^+^Endomucin^+^ cells but no tamoxifen-independent ZsGreen fluorescence is detected in representative Cre^-^ mice. (D) Flow cytometry analysis of adult BM cells from Cdh5-Cre^ERT2^(PAC)/ZsGreen (n = 6-8) and Cdh5-Cre^ERT2^(BAC)/ZsGreen mice (n = 10) shows that ZsGreen fluorescence identifies most ECs four weeks after tamoxifen administration but also tracks a small proportion of EC expressing tamoxifen-independent fluorescence in Cre^+^ but not Cre^-^ mice. (E) Percent CD45^+^ZsGreen^+^ cells of viable BM cells from Cre^-^ control (n=6), Cre^+^ control (peanut-oil treated, no tamoxifen; n= 8) and Cre^+^ tamoxifen-treated Cdh5-Cre^ERT2^(PAC) /ZsGreen (n=6) or Cdh5-Cre^ERT2^(BAC)/ZsGreen mice (n=9). Mice were 8 to 12 weeks old at the time of tamoxifen administration. (F) Percent ZsGreen^+^ cells of PBMC from Cre^-^ control, Cre^+^ control (peanut-oil treated) and Cre^+^ tamoxifen-treated Cdh5-Cre^ERT2^(PAC)/ZsGreen and Cdh5-Cre^ERT2^(BAC)/ZsGreen mice (n=9/group). Mice were 8 to 12 weeks old at the time of tamoxifen administration. (G–H) Representative flow cytometry gating of CD45⁺EGFP⁺ cells from BM and blood of Cdh5-CreERT2/mTmG mice (G, relates to Figure 1 C, D), and CD45⁺ZsGreen⁺ cells from BM and blood of Cdh5-Cre^ERT2^/ZsGreen mice (H, relates to Figure S1 E, F). (I) Representative confocal images showing a nucleated (DAPI^+^) BM ZsGreen^+^CD45^+^ cell in the BM from a Cdh5-Cre^ERT2^(PAC)/ZsGreen mouse treated with tamoxifen. (J) Percent ZsGreen^+^ cells of PBMC in individual Cdh5-Cre^ERT2^(BAC)/ZsGreen mice before or four weeks after tamoxifen administration. Each dot represents the results from 50-250µl blood/mouse. The lines link results from individual mice (n=15, 8-12 weeks old). (K) Percent B and T-lymphocytes, granulocytes, and monocytes among EGFP^+^ PBMC of Cre^+^ tamoxifen-treated Cdh5-Cre^ERT2^(PAC)/mTmG mice (n=15, 8-12 weeks old). (L) Percent EGFP^+^ cells in peripheral blood cell populations of Cre^+^ peanut oil-treated (n=9) and tamoxifen-treated (n=16) Cdh5-Cre^ERT2^(PAC)/mTmG mice (8-12 weeks old). Dots represent individual mice. Data are shown as mean ± SD. *p<0.05, **p<0.01, ***p < 0.001 by Student’s t test.

**Figure S2.**
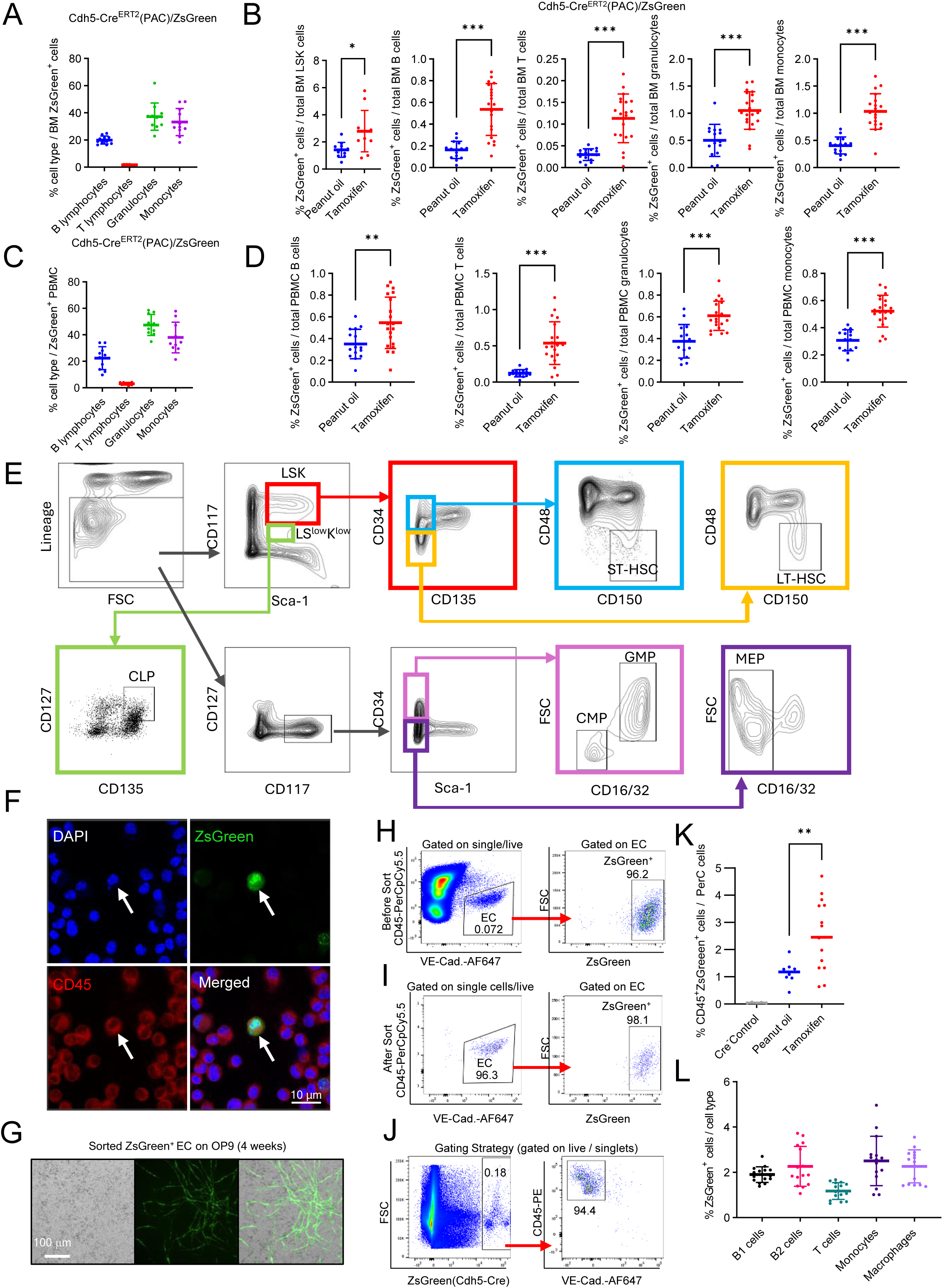
Characterization of tracked hematopoietic progenitors and mature cells in adult BM and peripheral blood of Cdh5-Cre reporter mice. Related to Figure 1, 2, or 3. (A and B) Percent B lymphocytes, T lymphocytes, granulocytes, and monocytes of all ZsGreen^+^ cells (A) and percent ZsGreen^+^ cells of total BM LSK, B and T-lymphocytes, granulocytes, and monocytes (B). Each dot reflects results from individual mice (1 femur plus 1 tibia combined, n=11); group means ± SD (error bars) are shown by the horizontal lines. (C and D) Percent B lymphocytes, T lymphocytes, granulocytes, and monocytes of all ZsGreen^+^ PBMC in Cdh5-Cre^ERT2^(PAC)/ZsGreen mice treated with tamoxifen (C, n=10) and percent ZsGreen^+^ cells of peripheral blood B lymphocytes, T lymphocytes, granulocytes, and monocytes in Cdh5-Cre^ERT2^(PAC)/ZsGreen mice (D) treated with peanut oil (n=15) or tamoxifen (n=20). (E) Gating strategy for identification of HSPC subsets in bone marrow. Representative flow cytometry plots show sequential gating of lineage negative (Lin^-^) Sca1^+^ cKit^+^ (LSK) cells into long-term hematopoietic stem cells (LT-HSC), short-term hematopoietic stem cells (ST-HSC), multipotent progenitors (MPP), common lymphoid progenitors (CLP), common myeloid progenitors (CMP), megakaryocyte-erythroid progenitors (MEP), and granulocyte-macrophage progenitors (GMP). (F) Representative cytospin image of floating and low-adherent cells from ex vivo culture of BM cells from a tamoxifen-induced Cdh5-Cre^ERT2^(PAC)/ZsGreen mouse. (G) Representative image (relates to Figure 2B) showing the appearance of sorted ZsGreen⁺ ECs after 4-week culture on OP9 monolayer. (H) Gating strategy for sorting VE-Cadherin^+^ ZsGreen^+^ ECs from the BM of Cdh5-Cre^ERT2^(PAC)/ZsGreen mice. (I) Purity analysis of sorted VE-Cadherin^+^ ZsGreen^+^ ECs. (J) Gating strategy used for detecting ZsGreen^+^CD45^+^ hematopoietic cells in WT C57Bl/6 recipients of ECs sorted from the BM of Cdh5-Cre^ERT2^(PAC)/ZsGreen donors. (K) Percent CD45^+^ZsGreen^+^ cells recovered from the peritoneal cavity (PerC) of Cre^+^Cdh5-Cre^ERT2^/ZsGreen mice treated with peanut oil (n=8) or tamoxifen (n=15). Cre^-^ mice (n=5). (L) Percent ZsGreen^+^ cells within PerC cell populations recovered from mice (n=15) under steady-state conditions. Cell type identification; B1 cells: CD19^+^, CD3^-^, CD45R(B220)^-^, CD5^+^, CD43^+^; B2 cells: CD19^+^, CD3^-^, CD45R(B220)^+^, CD5^-^, CD43^-^; T cells: CD3^+^, CD11b^-^; monocytes: CD11b^+^, CD19^-^, Ly6G^-^, Ly6C^high^; neutrophils: CD11b^+^, CD19^-^, Ly6G^+^, Ly6C^low^; and macrophages: CD11b^+^, CD19^-^, F4/80^+^. Dots represent individual mice. Data are shown as mean ± SD. *p<0.05, **p<0.01, ***p < 0.001 by Student’s t test.

**Figure S3.**
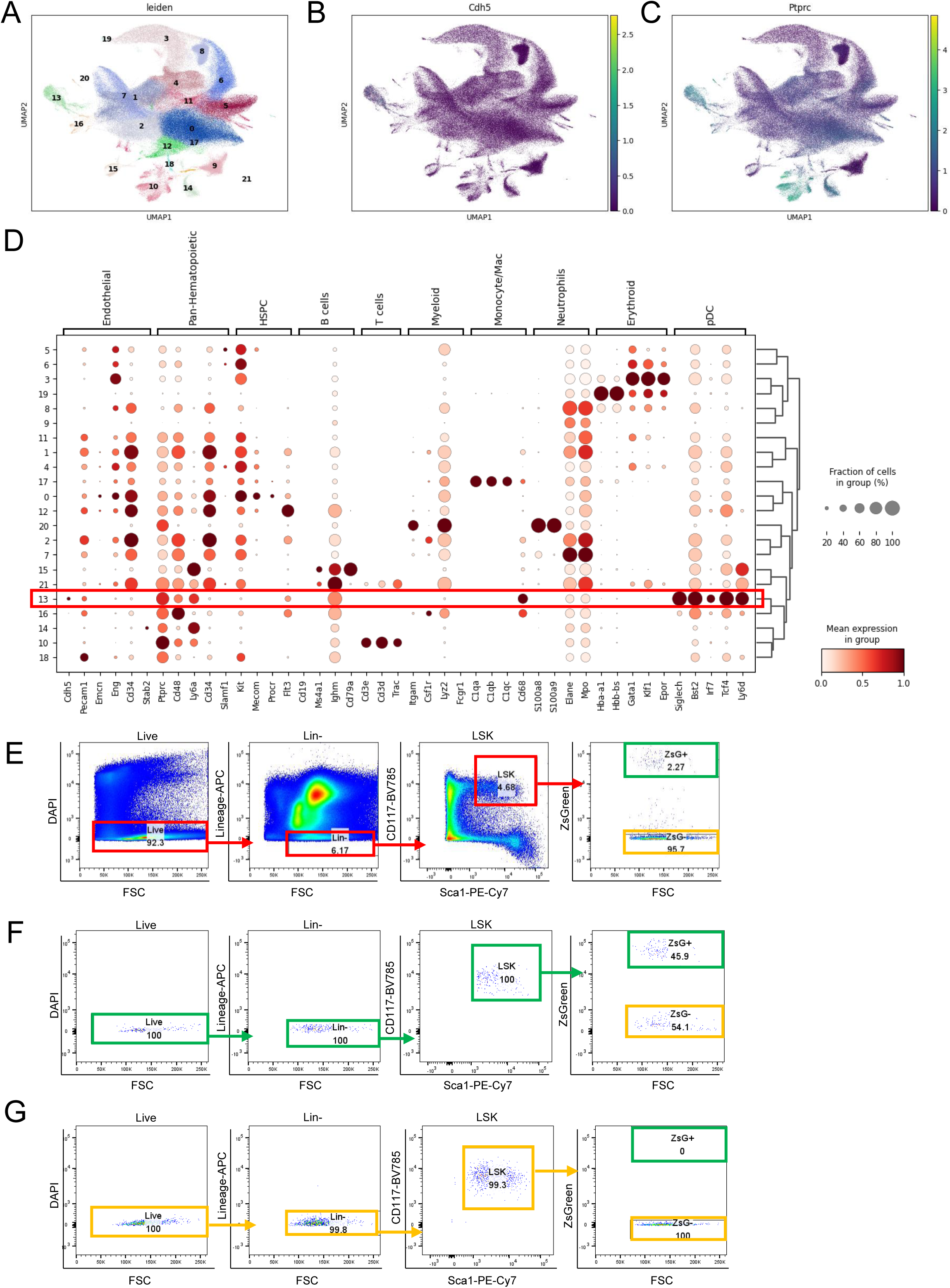
BM pDCs, but not other BM hematopoietic cells, express *Cdh5* and *Ptprc*, encoding CD45. Related to Figure 4. (A) UMAP plot showing unsupervised clustering of sc transcriptomic data from a public dataset of mouse BM hematopoietic cells. (B and C) UMAP plots displaying expression of *Cdh5* (B) and *Ptprc* (encoding CD45, C) in the dataset shown in (A). (D) Dot plot illustrating the expression of selected marker genes across clusters shown in (A). The red rectangle highlights gene expression by cluster 13 cells, identifying pDCs. Dot size represents the percentage of cells expressing the gene within each cluster, and color intensity reflects the mean expression level. (E) Gating strategy for selecting ZsGreen^-^ and ZsGreen^+^ LSK progenitors. (F and G) Analysis of purity of sorted LSK populations enriched for ZsGreen^+^ cells (F) and depleted of ZsGreen^+^ cells (G) from the BM of Cdh5-Cre^ERT2^(PAC)/ZsGreen mice (not treated with tamoxifen).

**Figure S4.**
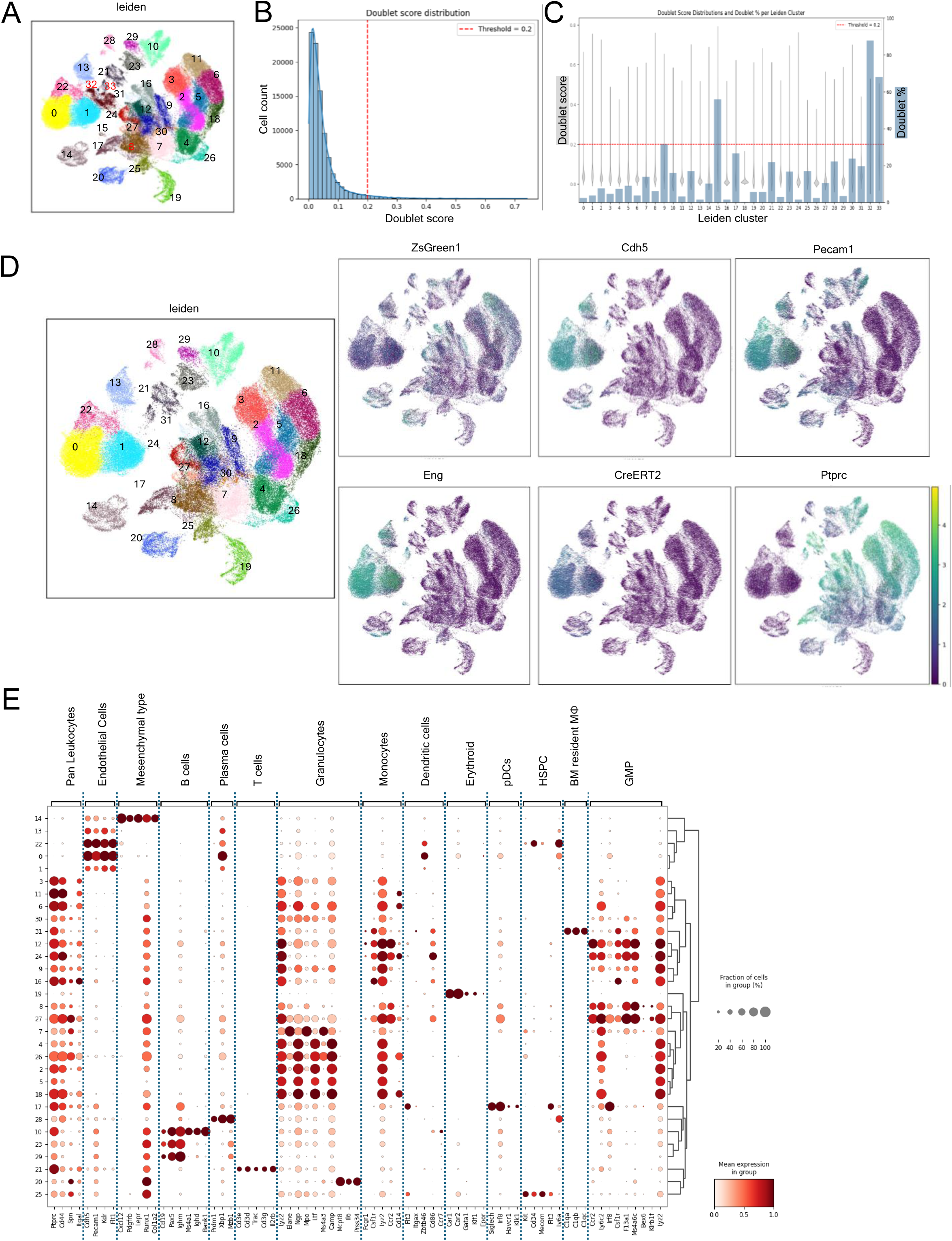
Single-cell RNA-seq analysis of BM ZsGreen^+^ cells from tamoxifen-treated Cdh5-Cre/ZsGreen/Polylox mice. Related to Figure 5. (A) UMAP plot showing unsupervised clustering of sc RNA-seq data, identifying 34 distinct cell clusters within BM ZsGreen^+^ cells. (B) Histogram of doublet score distribution. A threshold of 0.2 was applied to match the expected doublet rate from 10x Genomics Chromium GEM-X chips. (C) Doublet score distribution (gray violin plots, left Y-axis) and corresponding doublet percentages (blue bars, right Y-axis) across Leiden clusters. (D) UMAP plots of clusters after doublet removal, and expression of *ZsGreen1*, *Cdh5*, *Pecam1*, *Eng*, *CreERT2* and *Ptprc* (CD45). (E) Dot plot showing expression of selected marker genes across Leiden clusters identified in (A). Dot size indicates the proportion of cells expressing the gene; color intensity reflects the average expression level of each cluster.

**Figure S5.**
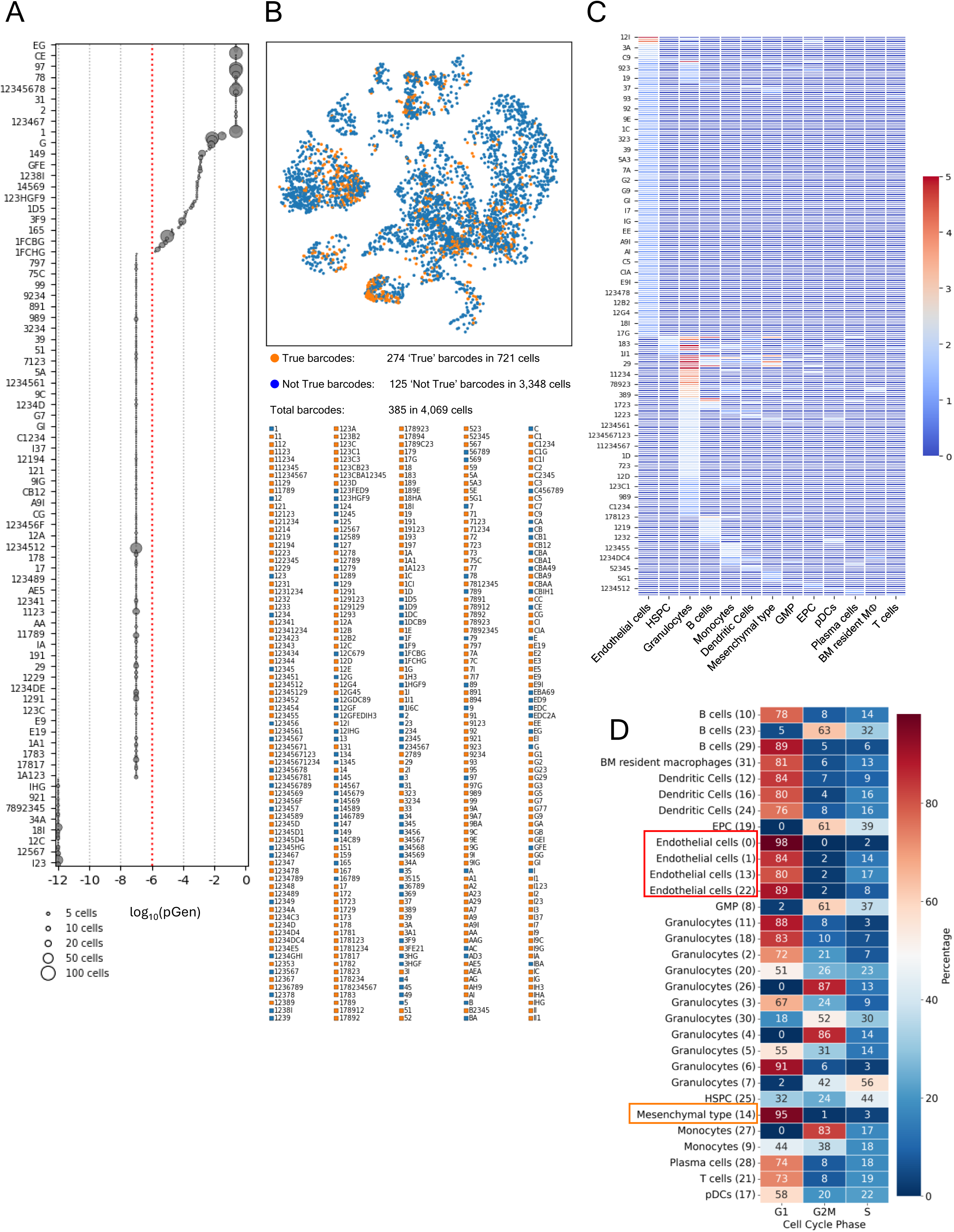
Identification and distribution of ‘True’ Polylox barcodes across cell types. Related to Figure 5. (A) Bubble plot showing individual Polylox barcodes plotted against their corresponding pGen values (log₁₀ scale). The y-axis indicates representative barcodes (one label is shown for every five barcodes). Bubble size reflects the number of cells harboring each barcode. The red dashed line denotes the log₁₀(pGen) = −6 cutoff, which was used to define true barcodes retained for downstream analyses. (B) UMAP plot showing the distribution of ‘True’ Polylox barcodes across the 34 Leiden-defined clusters identified in Figure S4A. Cells containing ‘True’ barcodes (n = 721) are shown in orange; cells with ‘Not True’ barcodes (n = 3,348) are shown in blue. A total of 388 barcodes were detected, including 274 ‘True’ (orange) and 125 ‘Not True’ (blue), as listed below the UMAP plot. (C) Heatmap displaying the distribution and abundance of ‘True’ Polylox barcodes across annotated cell types. Each row corresponds to a unique barcode (1 out of every 5 barcodes shown); color intensity represents the number of cells carrying that barcode within each listed cell type. (D) Heatmap showing cell cycle phase distribution (G1, G2/M, S) across Leiden clusters identified in Figure S4A. Color intensity and numerical values represent the percentage of cells in each phase within the indicated cell type.

**Figure S6.**
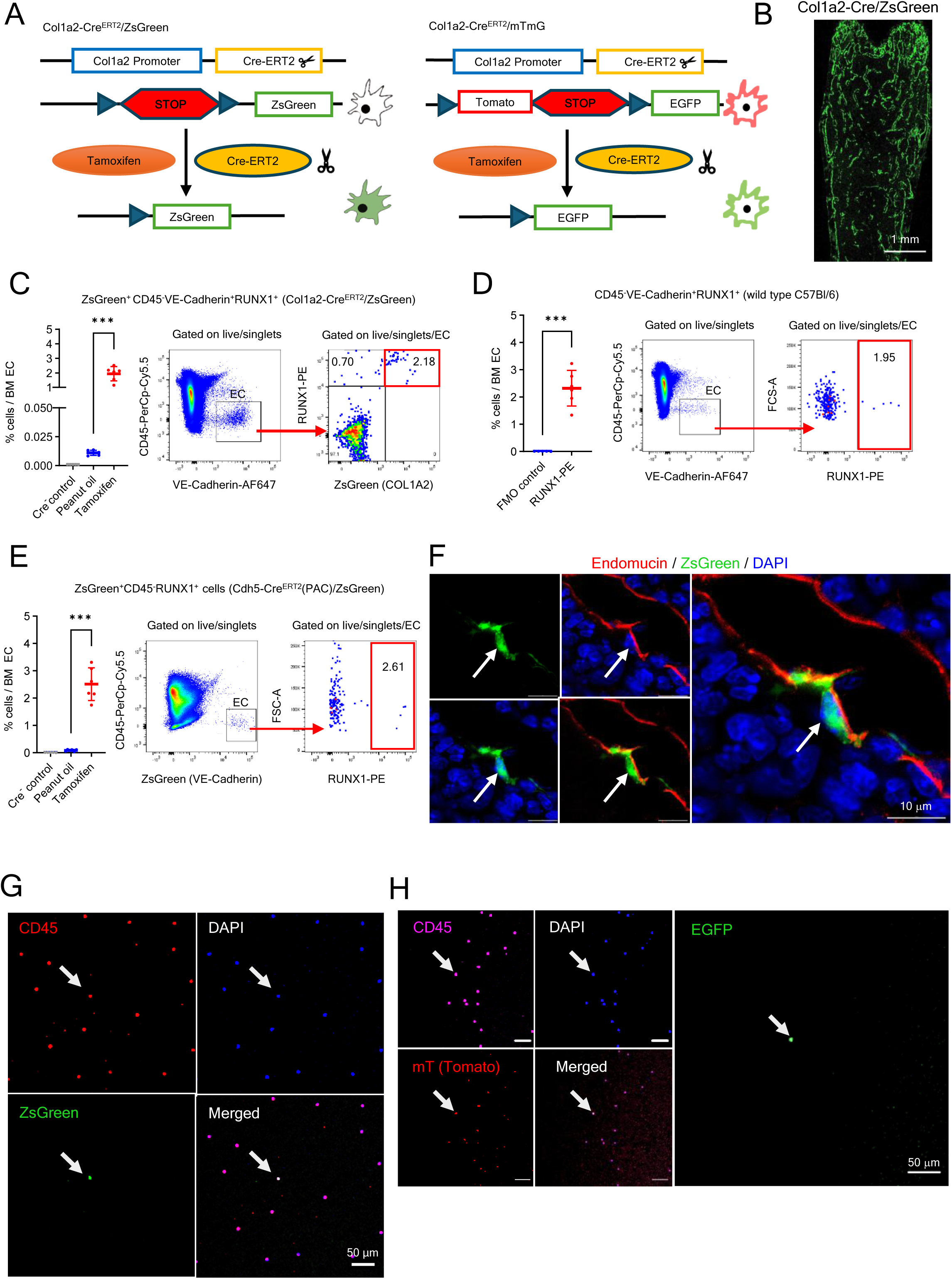
Characterization of Col1a2-tracked cell populations in BM and blood. Related to Figure 7. (A) Schematic representation of the Col1a2 tracking lines. (B) Representative confocal image of a BM section from a tamoxifen-treated Col1a2-Cre^ERT2^/ZsGreen mouse, showing widespread distribution of ZsGreen⁺ cells. (C to E) Flow cytometric identification of RUNX1^+^VE-Cadherin^+^CD45^-^ ECs in the BM of peanut oil-treated ( n=6) and tamoxifen-treated (n=6) Col1a2-Cre^ERT2^/ZsGreen adult mice; Cre^-^mice (n=5) (C); WT C57Bl/6 mice (n= 6) and Fluorescence Minus One (FMO) control (n=5) (D); a^nd C^dh5-Cre^ERT2^(PAC)/ZsGreen mice treated with peanut oil (n=6) or tamoxifen (n=6); Cre^-^ mice (n=5)(E). Left panels: quantification of cells identified by the indicated gates as a percentage of total BM ECs; each dot represents one mouse (1 femur + 1 tibia). Middle and right panels: representative gating strategies. (F) Representative confocal microscopy image of a BM section from a tamoxifen-treated Col1a2-Cre^ERT2^/ZsGreen adult mouse showing a ZsGreen^+^ Endomucin^+^ cell lining a vascular structure (white arrows). (G) Representative confocal image of a blood smear from a Col1a2-Cre^ERT2^/ZsGreen mouse treated with tamoxifen showing the presence of a nucleate CD45^+^ cell tracked by ZsGreen/Col1a2 fluorescence (pointed by the arrow). (H) Representative confocal image of a blood smear from a Col1a2-Cre^ERT2^/mTmG mouse treated with tamoxifen showing the presence of a nucleated CD45^+^ cell tracked by EGFP/Col1a2 fluorescence (pointed by the arrow). Dots represent individual mice. Data are shown as mean ± SD. ***p < 0.001 by Student’s t test.

**Figure S7.**
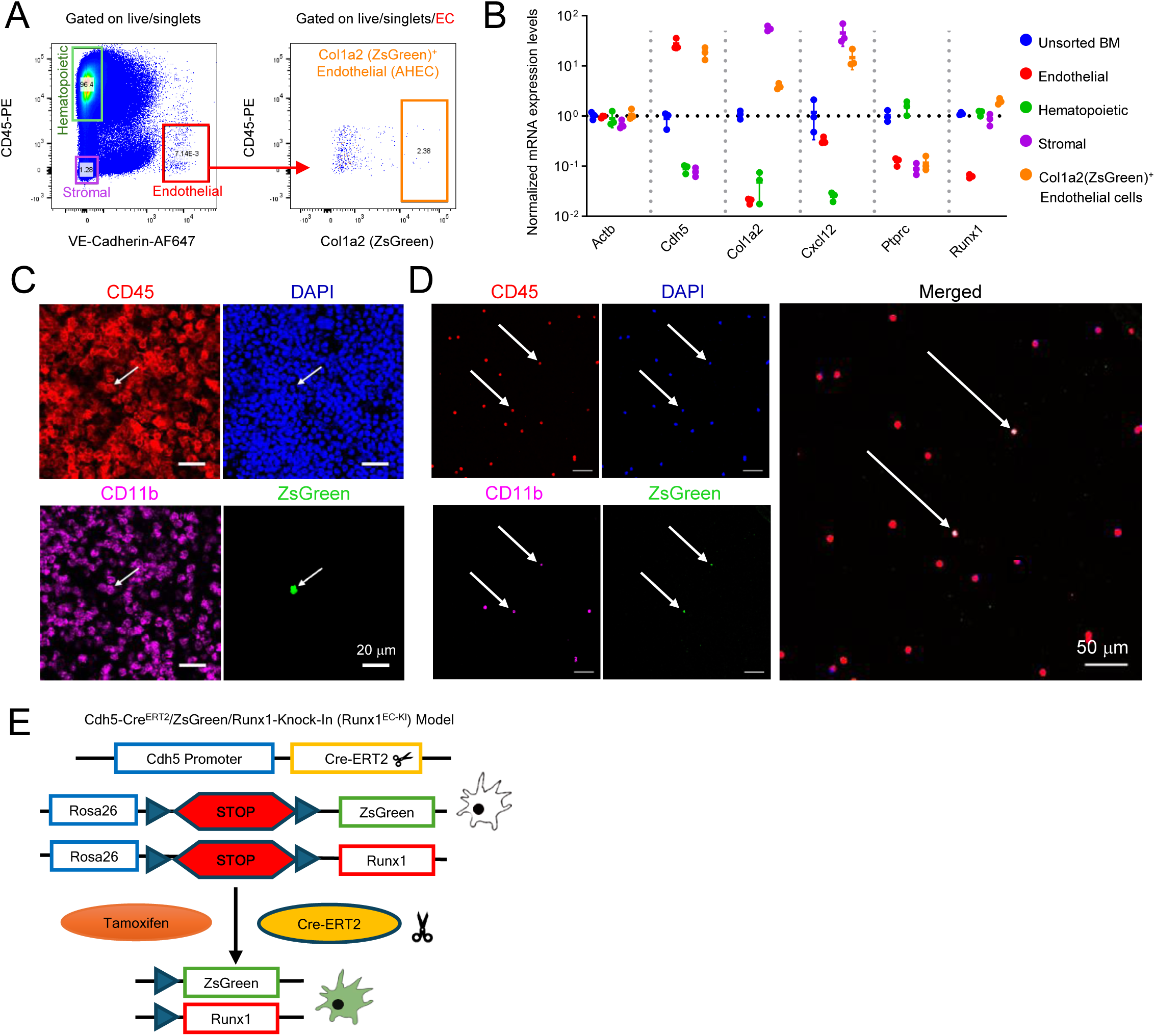
Analysis and hemogenic potential of Col1a2-tracked adult BM ECs. Related to Figure 7. (A) Representative FACS gating strategy used to isolate BM hematopoietic cells, ECs, stromal cells, and Col1a2-tracked ECs from tamoxifen-treated Col1a2-Cre^ERT2^/ZsGreen mice. (B) Gene expression profiling of unsorted BM and sorted BM populations defined in (A). Results from qRT-PCR are normalized by *Gapdh* and unsorted BM. Dots reflect experimental triplicates. (C) Representative confocal image of a BM section from a transplant recipient showing a CD45⁺CD11b⁺ZsGreen⁺ cell (arrow), indicating hematopoietic derivation from transplanted ZsGreen⁺VE-Cadherin⁺CD45⁻ (Col1a2⁺) cells. (D) Representative image of a blood smear from a WT recipient mouse transplanted with ZsGreen (Col1a2)^+^ BM ECs from Col1a2-Cre^ERT2^/ZsGreen mice. ZsGreen tracked cells are pointed by the arrows. (E) Schematic diagram of the Cdh5-Cre^ERT2^/ZsGreen/Runx1-Knock-in (Runx1^EC-KI^) mouse line used to trace endothelial cells with Runx1 expression induced upon tamoxifen treatment.

## Supplemental Method

### Polylox single-cell lineage tracing and barcode analysis

Polylox single-cell lineage tracing experiments were performed essentially as described previously (Pei et al., 2020), with modifications detailed below.

### Mouse treatment and sample collection

Cdh5-CreERT2/ZsGreen/PolyloxExpress mice were treated with tamoxifen at 10 weeks of age and harvested at 16 weeks of age (approximately 6 weeks post induction). Bone marrow (BM) was collected from three mice (n = 3, 4 long bones from each mouse), separately processed for downstream analyses.

### Cell sorting and population mixing

BM cells were enriched for ZsGreen⁺ cells by fluorescence-activated cell sorting (FACS). The sorter was set up with 100-micron nozzle, 20psi. Sample rate was set at ∼6,000 EPS. Sorting purity was set as 4-way-purity.

Two populations were isolated at >99% purity:

1. ZsGreen⁺ VE-Cadherin⁺ Endomucin⁺ endothelial cells (ECs), and
2. ZsGreen⁺ VE-Cadherin⁻ Endomucin⁻ CD45⁺ hematopoietic cells.

The two sorted populations were mixed at a 1:1 ratio prior to single-cell encapsulation to allow direct comparison of barcode sharing across endothelial and hematopoietic compartments. The six samples from three mice were loaded separately onto 6 different lanes of a 10x GEM-X and processed as individual samples moving forward.

### Single-cell capture and sequencing

A total of 147,446 cells were loaded onto the 10x Genomics Chromium platform, of which 93,553 cells were successfully processed and recovered for single-cell RNA sequencing. Reverse transcription and cDNA amplification were performed according to the manufacturer’s protocols. Amplified cDNA was split into two aliquots for parallel transcriptome library preparation and Polylox barcode enrichment.

### Transcriptome library preparation and sequencing

For transcriptome analysis, 25% (10 μL) of the amplified 10x cDNA library was fragmented and processed using Chromium Single Cell 3′ Reagent Kits (v3 and v4). Libraries were sequenced on an Illumina NextSeq 2000 using P3/P4 reagents (28 bp read 1 + 74 bp read 2).

### Polylox barcode amplification and long-read sequencing

Polylox barcodes were enriched from 5–10 ng of amplified 10x cDNA by nested PCR. First-round PCR was performed using primers annealing to the 5′ end of the Polylox cassette and the 10x adaptor sequence, followed by a second round of amplification using internal Polylox primers. PCR products were purified using AMPure beads and used to generate long-read amplicon libraries, which were sequenced on a PacBio Sequel II platform using SMRTbell library preparation (v3.0).

### Barcode recovery and integration with transcriptomes

Polylox barcodes were extracted using a Snakemake workflow with custom Python scripts (https://github.com/CCRSF-IFX/SF_Polylox-BC). Barcode sequences were linked to single-cell transcriptomes through shared 10x cell indices. A custom mm10 reference genome including ZsGreen1 was used for transcriptome alignment.

### Barcode filtering and definition of true barcodes

Barcode generation probabilities (pGen) were calculated using the published Polylox MATLAB pipeline (https://github.com/hoefer-lab/polylox). “True” barcodes were defined as barcodes with pGen < 1 × 10⁻^6^. Using this threshold, 296 true barcodes were retained for downstream lineage analysis.

### Clone size and abundance analysis

Clone sizes and barcode abundances across endothelial, hematopoietic progenitor, mesenchymal-type, and mature blood cell populations were quantified based on the number of cells sharing the same true barcode. These data are visualized as heatmaps (Figure 5C–E) and as intersection plots summarizing barcode sharing across cell types (Figure 5F).

### Quality control and downstream analysis

Doublets were identified using Scrublet, with expected doublet rates estimated based on cell loading numbers. Batch correction was performed using BBKNN. Cell type annotations were assigned based on canonical marker expression and supported by decoupleR, scANVI (ImmGen reference), and CellTypist. All single-cell analyses were performed using Scanpy.

| cell_type | Total cells | Cells with barcodes (pGen < 1) | Cells with true barcodes (pGen < 1e-6) | Cells with not true barcodes (1e-6 <= pGen < 1) |
| --- | --- | --- | --- | --- |
| B cells | 5977 | 344 | 73 | 271 |
| BM resident macrophages | 547 | 18 | 5 | 13 |
| Dendritic Cells | 6471 | 235 | 26 | 209 |
| Doublets | 2607 | 87 | 17 | 70 |
| Endothelial cells | 21168 | 828 | 169 | 659 |
| Erythroblasts and erythroid precursors | 1809 | 78 | 13 | 65 |
| GMP | 3980 | 121 | 11 | 110 |
| Granulocytes | 37931 | 1674 | 317 | 1357 |
| HSPC | 1257 | 71 | 14 | 57 |
| Mesenchymal type | 2544 | 115 | 31 | 84 |
| Monocytes | 4840 | 281 | 40 | 241 |
| Plasma cells | 824 | 49 | 8 | 41 |
| T cells | 1580 | 52 | 3 | 49 |
| pDCs | 2018 | 119 | 11 | 108 |

